# Is it the same strain? Defining genomic epidemiology thresholds tailored to individual outbreaks

**DOI:** 10.1101/2022.02.15.480545

**Authors:** Audrey Duval, Lulla Opatowski, Sylvain Brisse

## Abstract

**Background:** Epidemiological surveillance relies on microbial strain typing, which defines genomic relatedness among isolates to identify case clusters and their potential sources. No consensus exists on the choice of thresholds of genomic relatedness to define clusters. While *a priori* defined thresholds are often applied, outbreak-specific features such as pathogen mutation rate and duration of source contamination should be considered.

**Methods:** We developed a forward model of bacterial evolution to simulate mutation within a population diversifying at a specific mutation rate, with specific outbreak duration and sample isolation dates. Based on the resulting expected distribution of genetic distances we define a threshold beyond which isolates are considered as not part of the outbreak. We additionally embedded the model into a Markov Chain Monte Carlo inference framework to estimate, from data including sampling dates or isolates genetic variation, the most credible mutation rate or time since source contamination.

**Findings:** A simulation study validated the model over realistic durations and mutation rates. When applied to 16 published datasets describing foodborne outbreaks, our framework consistently identified outliers. Appropriate thresholds for grouping cases were obtained for 14 outbreaks. For the remaining two outbreaks, re-estimation of the duration of outbreak lead to updated threshold values and was more likely, given our model, to result in the observed genetic distances.

**Interpretation:** We propose an evolutionary approach to the ‘single strain’ conundrum by defining the genetic threshold based on individual outbreak properties. The framework provides an informed estimation of the likelihood of a cluster given the samples epidemiological and microbiological context. This forward model, applicable to foodborne or environmental-source single point case clusters or outbreaks, will be useful for epidemiological surveillance and to guide control measures.

**Funding:** This work was supported financially by the MedVetKlebs project, a component of European Joint Programme One Health EJP, which has received funding from the European Union’s Horizon 2020 research and innovation programme under Grant Agreement No 773830. The funders had no role in study design, data collection and analysis, decision to publish, or preparation of the manuscript.

**Research in context:** *Evidence before this study:* We searched PubMed for studies published between database inception and April 3, 2021, with the term (threshold OR cut-off OR genetic relatedness) AND (outbreak) AND (cgMLST OR wgMLST OR SNPs) AND (microbial OR bacteria OR bacterial OR pathogen). We found 222 related articles. Most studies define a fixed SNP threshold that relate outbreak strains based on previous observations. One original study identifies outbreak clusters based on transmission events. However, it relies on strong assumptions about molecular clock and transmission processes.

*Added value of this study:* Our study describes a new method based on a forward Wright-Fisher model to find the most credible genetic distance threshold. This method is fast and simple to use with only few assumptions, informed by outbreak duration and pathogen mutation rate. By using SNP or cgMLST pairwise distances and sample collection dates of the outbreak of interest, the algorithm provides context-based guidance to separate outbreak strains from outliers.

*Implications of all the available evidence:* The fast and easy method developed here enables to move away from *a priori* defined thresholds. Defining clusters more accurately based on the specific features of outbreaks, and the ability to estimate outbreak duration, will provide the needed precision for epidemiological surveillance and should contribute to leverage molecular epidemiology data more efficiently for the purpose of uncovering contamination sources.

*Data Availability Statement:* All data and code used for this manuscript is available online at https://gitlab.pasteur.fr/BEBP.

## Introduction

Outbreaks of infections caused by the exposure to a unique source are the particular focus of surveillance and infection control strategies. The rapid identification of the source can lead to immediate public health benefits and is therefore critical. In the simplest cases, a single strain of infectious agent contaminates the source and subsequently causes infections (referred to as a ‘clonal outbreak’). This is often the case for contaminated food, water or environmental sources that are under strong regulatory measures and typically uncontaminated. Surveillance systems were therefore put in place, *e*.*g*., for food-borne pathogens such as *Salmonella* or *Listeria monocytogenes*, based on a collect-genotype-compare strategy [1,2]. This strategy, dubbed ‘reverse epidemiology’ [3], forms the basis of surveillance systems for foodborne pathogens, such as PulseNet [1]. Molecular surveillance (‘genetic fingerprinting’) enables the detection of nearly identical infectious agent isolates and may trigger epidemiological investigations. These include the search for case-associated risk factors as well as microbiological analyses of suspected sources, and may lead to infection control measures that can prevent further infections.

Distinguishing case cluster isolates from sporadic ones has been the ‘Holy Grail’ of molecular epidemiological surveillance. However, the identification of single-strain clusters of infections is confounded by a background of sporadic cases caused by exposure to unrelated sources. Defining ‘a single strain’ typically involves the use of a threshold of genetic distance, which discriminates between isolates that are related or not to the event. The literature is ripe with attempts to define such thresholds [4]. In the whole-genome sequencing (WGS) era, thresholds were refined compared to pre-genomic methods such as PFGE [5–10]. Usually, threshold definition is based on the variability observed within previously well-characterised outbreaks, an approach rooted in the epidemiological concordance principle [11]. However, interpretation of molecular data for strain definition is far from being consensual [5,12,13].

From an evolutionary biology point of view, infectious agents that are present as contaminants of an initially sterile source can be considered as subpopulations of individuals that have evolved from a single common ancestor (the original strain) since some time (the duration of contamination). Major factors expected to influence the genetic distances among sampled individuals (isolates) include: i) the duration of strain persistence in the contaminated source prior to infections; ii) the evolutionary rate of the pathogen genomic markers; iii) the sampling dates. On the other hand, the genetic distance to the closest observed isolate unrelated by source will be determined by which genomes were sampled outside the contamination event. All these parameters considered, the quest for a unique threshold applicable to all outbreaks is deemed to fail. Instead, using outbreak-specific thresholds defined based on their context-informed expected diversity is likely to represent a more successful strategy. Attempts to ground threshold definition in evolutionary biology are recent and used the coalescent model [6], transmission models [14] and Bayesian MRCA models [15,16].

The aim of this work was the development of a novel model to define the most credible genetic distance cut-offs for single strain outbreaks from a contaminated source, by simulating the accumulation of mutations using specific outbreak parameters.

## Methods

### Evolutionary model and definition of the outbreak genetic distance threshold

We define an outbreak (or cluster of cases) as a group of infection cases caused by a single strain (‘monoclonal’), excluding co-occurring cases caused by genetically unrelated strains (*i*.*e*., from other sources). In the case where two or more genetically unrelated strains co-contaminate the source of the outbreak, they should be analysed separately with this framework.

Our evolutionary formalization (**Figure 1A)** is based on a Wright-Fisher forward model of haploid infectious agent evolution [17,18] with constant population size. The simulation is initialised with a homogeneous population of an infectious agent characterised by five properties: i) *L*, the genome length (base pairs, bp) or the average length of genes of multilocus sequence typing [MLST] approaches; ii) *g*, the number of genes; iii) *μ*, the number of substitutions per site per year; iv) *D*, the duration (in days) of the outbreak, defined as the time elapsed between the initial contamination of the source, and the sampling date of the last isolate; and v) *S*_*d*_, the set of sampling dates of isolates, which is defined either directly from the source sampling dates or from the date of sampling of infections, in which case the incubation time and within-patient evolution is neglected. Substitutions are introduced at each time step in individuals sampled with replacement according to a uniform distribution (Poisson distribution with parameter *λ*). A distribution of pairwise genetic distances is generated on these sampled individuals, and the genetic threshold value is defined from this distribution. Details of the model are provided in the **Supplementary Appendix**.

**Figure 1.**
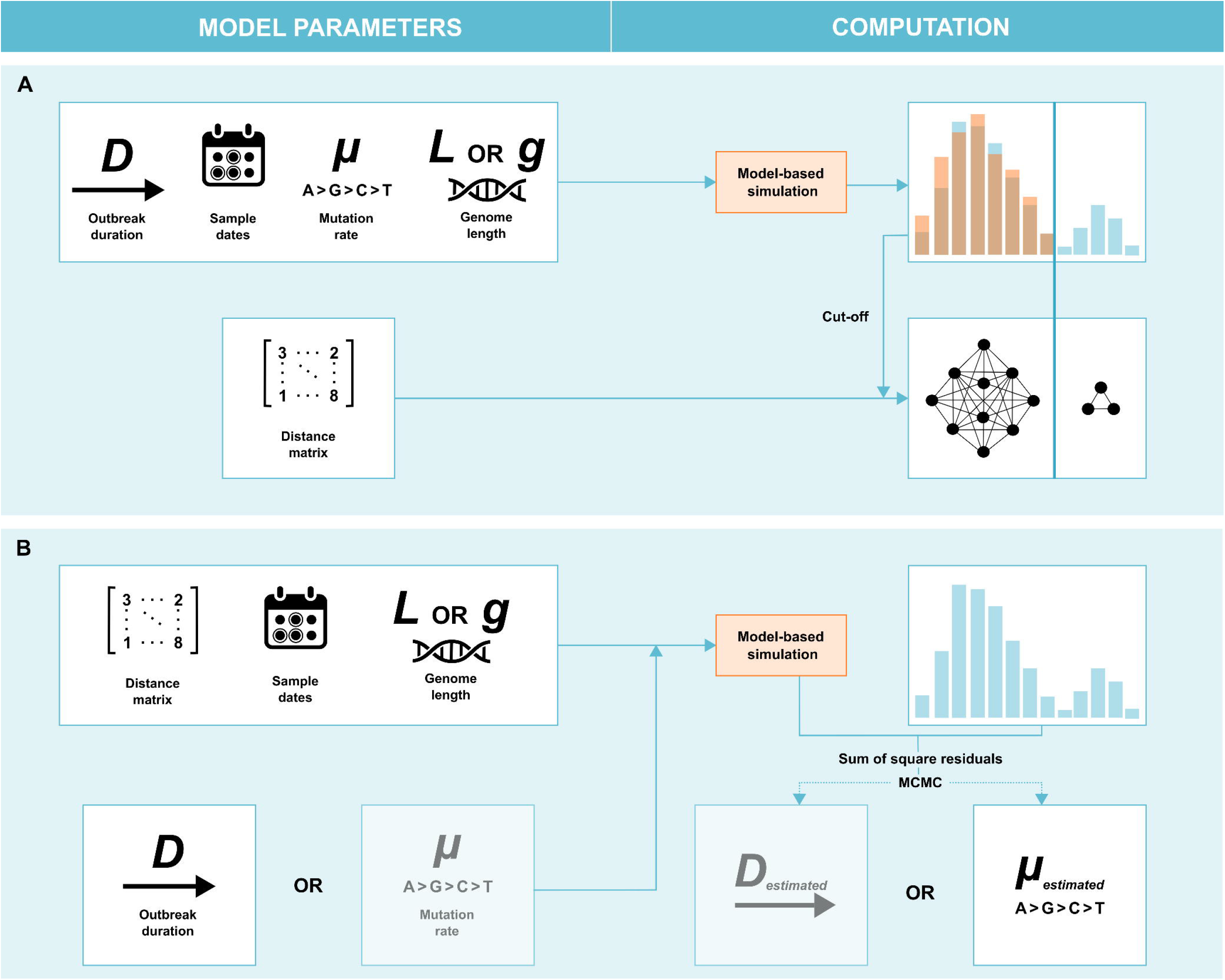
Description of the framework. **A**. Left, threshold computation inputs: genetic distance matrix *M*, duration of outbreak *D*, set of sample dates *S*_*d*_, number of substitutions per site per year *μ*, and sequence length *L* (if based on nucleotide sites) or number of genes *g* (if based on a gene-by-gene approach). Right, model-based simulation: the algorithm is initialized with a homogenous population of individuals. At each time step, substitutions are drawn from a Poisson distribution, until *D* is reached. Samples are drawn randomly at the different observed sampling dates. A genetic threshold is defined using *e*.*g*., the 99^th^ percentile of the distribution, and clusters of isolates are derived by single linkage clustering, leading to rule-out non-outbreak isolates. **B**. Left, the same model is used to estimate *D* or *μ* using MCMC, based on the following inputs: the genetic distance matrix; the sampling dates; the sequence length; and either *μ* or *D* (depending on which one is estimated).

### Analysis of published outbreak datasets

We reviewed available published outbreak datasets from the literature and analysed the 16 datasets listed in **Table 1** [6,19–24] using our modelling framework. Inclusion criteria were i) foodborne outbreak; ii) the availability of whole genome sequence data and iii) availability of collection dates of isolates. The 16 outbreaks are described in more details in the **supplementary appendix**. We estimated *D* based on evidence provided in the original publications on these outbreaks. We also used previously estimated *μ* and *g* for the corresponding infectious agent from literature (**Table 1**). We labelled *D* and *μ* values taken from the literature as *D*_*lit*_ and *μ*_*lit*_, whereas those derived from our Markov Chain Monte Carlo (MCMC) estimation (see below) were labelled as *D*_*estimated*_ and *μ*_*estimated*._

**Table 1.**
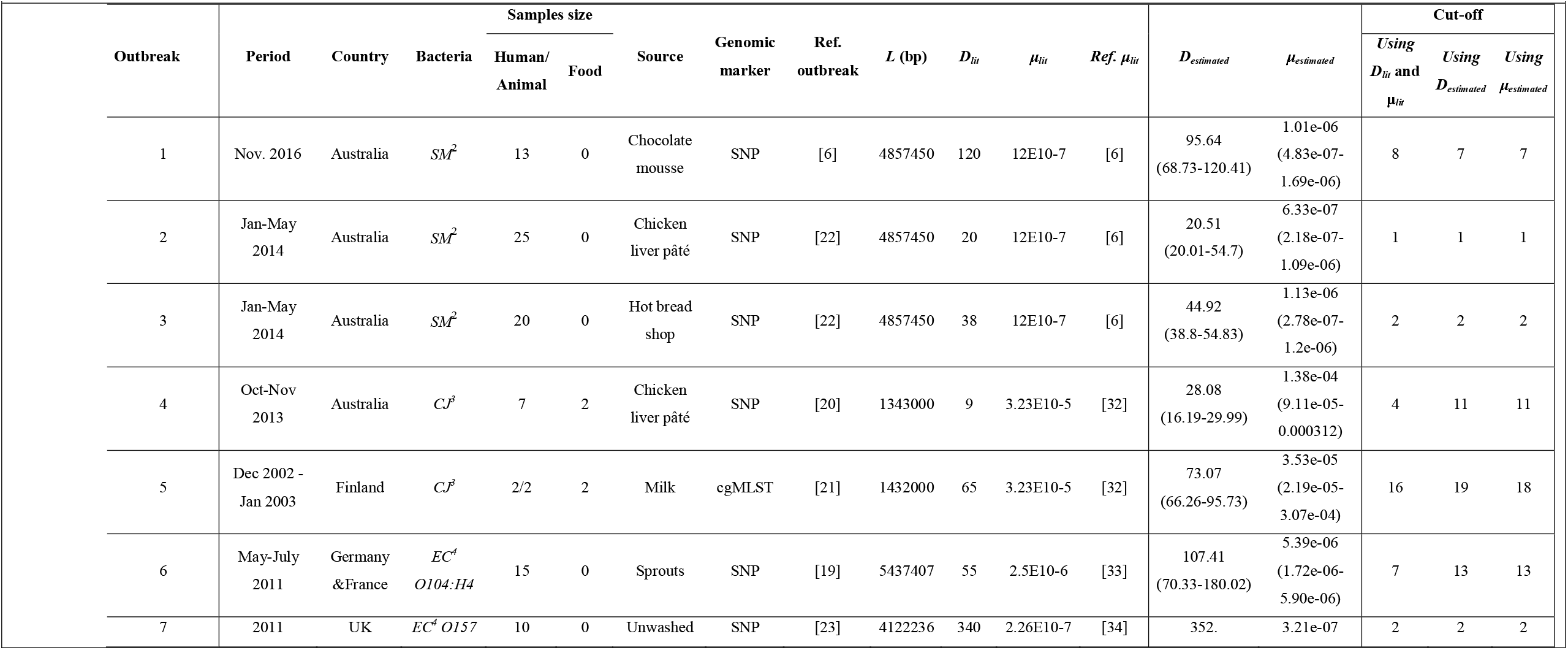
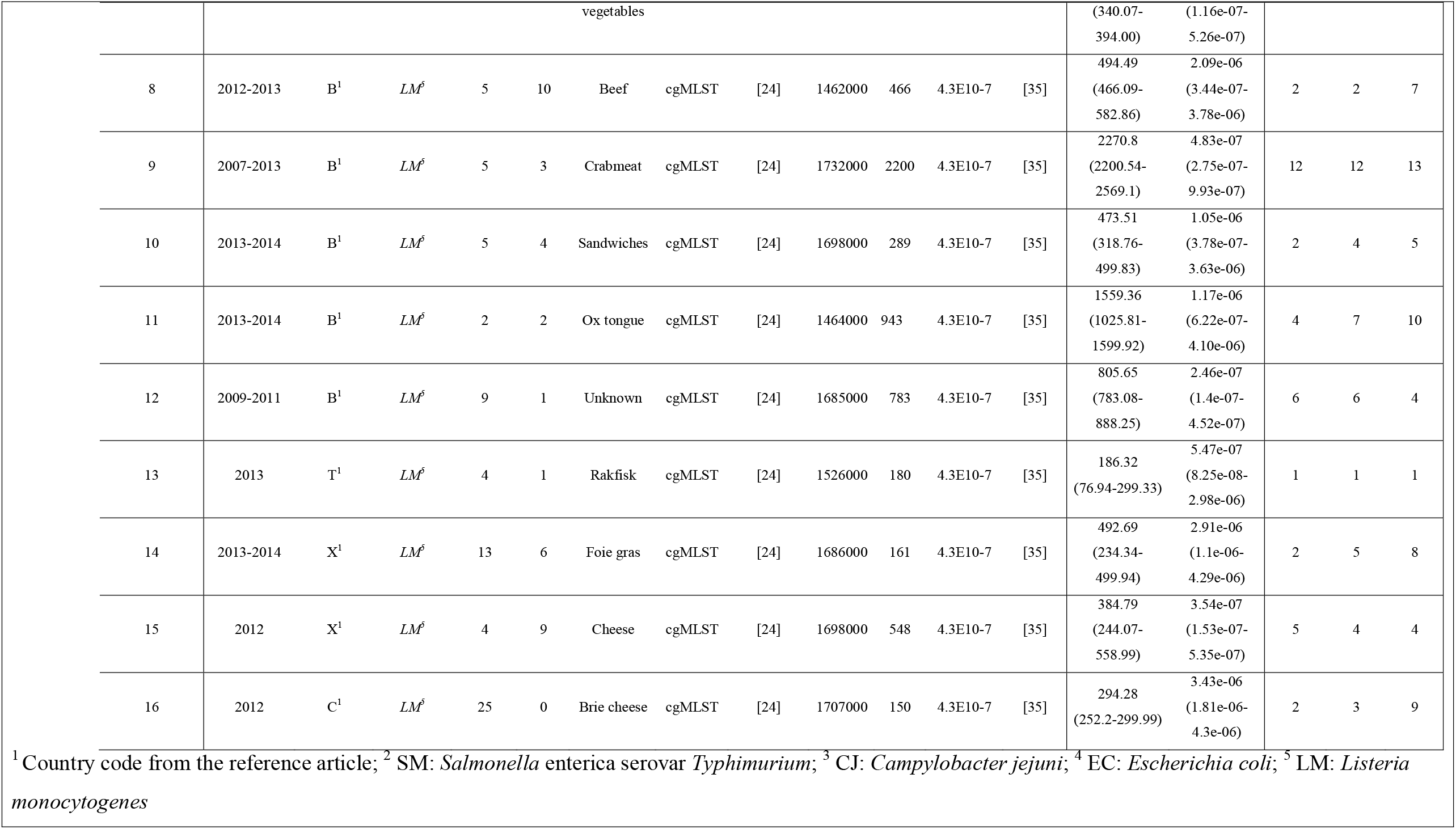
Analysis of 16 outbreaks from literature. Genome length *L* is given in base pairs (bp), for outbreaks 5 and 8 to 16 the genome length was given by the number of loci *g* multiply by the average gene length. *D*_*lit*_ correspond to the duration of outbreak in days deduced from the published articles and *μ*_*lit*_ is the number of mutations per site per year found in the published article related or found in the literature in general. For each *μ*_*lit*_ value, the reference used is shown. *D*_*estimated*_ (in days) and *μ*_*estimated*_ were estimated based on 3 MCMC chains; the associated 95% HPD for all the outbreaks as well as the corresponding genetic threshold is shown. Each threshold results from the 99^th^ percentile of a pairwise differences distribution from 100 outbreak simulations.

### Statistical analyses, simulation studies and statistical framework

#### Model assessment

To assess the capacity of the model to adequately tell apart outbreak isolates from non-outbreak isolates, we used synthetic datasets generated with different parameters values. We applied our framework to a series of 171 simulated outbreaks generated with 19 different values of *D* each combined with 9 values of *μ* and including simulated sporadic isolates (**Table S1** in the supplementary appendix). For each of them, we assessed the global sensitivity (*Se*) and specificity (*Sp*) of the framework. Details are provided in the **Supplementary Appendix**.

#### Parameters estimation

Our model was embedded into a Bayesian inference statistical framework to enable estimation of either the duration (*D*) or the substitution rate (*μ*) of studied outbreaks (**Figure 1B; Supplementary appendix**). Simulated outbreaks were used to assess the ability of the model to estimate *D* and *μ*, and their impact on the genetic threshold estimation. We used the Kolmogorov-Smirnoff test statistic (noted *D*_*KS*_) to compare real distributions with simulated distributions as a goodness of fit indicator. Details on the inference framework are provided in the **Supplementary Appendix**.

## Results

### Analysis of simulated outbreaks: accuracy of outbreak delineation and of parameters estimation

To test the ability of the framework in distinguishing between outbreak and non-outbreak cases, we generated synthetic outbreaks from different combinations of *D* and *μ* (**Table S1 in the supplementary appendix**). **Figure 2** shows the specificity *Sp* and sensitivity *Se* according to *μ*. *Sp* was poor with low *μ* values, especially when *R*_*d*_ (the ratio of evolution duration between outbreak and non-outbreak genomes) was small (**Figure 2A**). In contrast, *Se* was always high (more than 99%, **Figure 2B**), irrespective of the parameter’s combinations. We observed that the higher *R*_*d*_ and *μ* were, the lower this 95% *Sp D*-value threshold was (**Figure 2C**).

**Figure 2.**
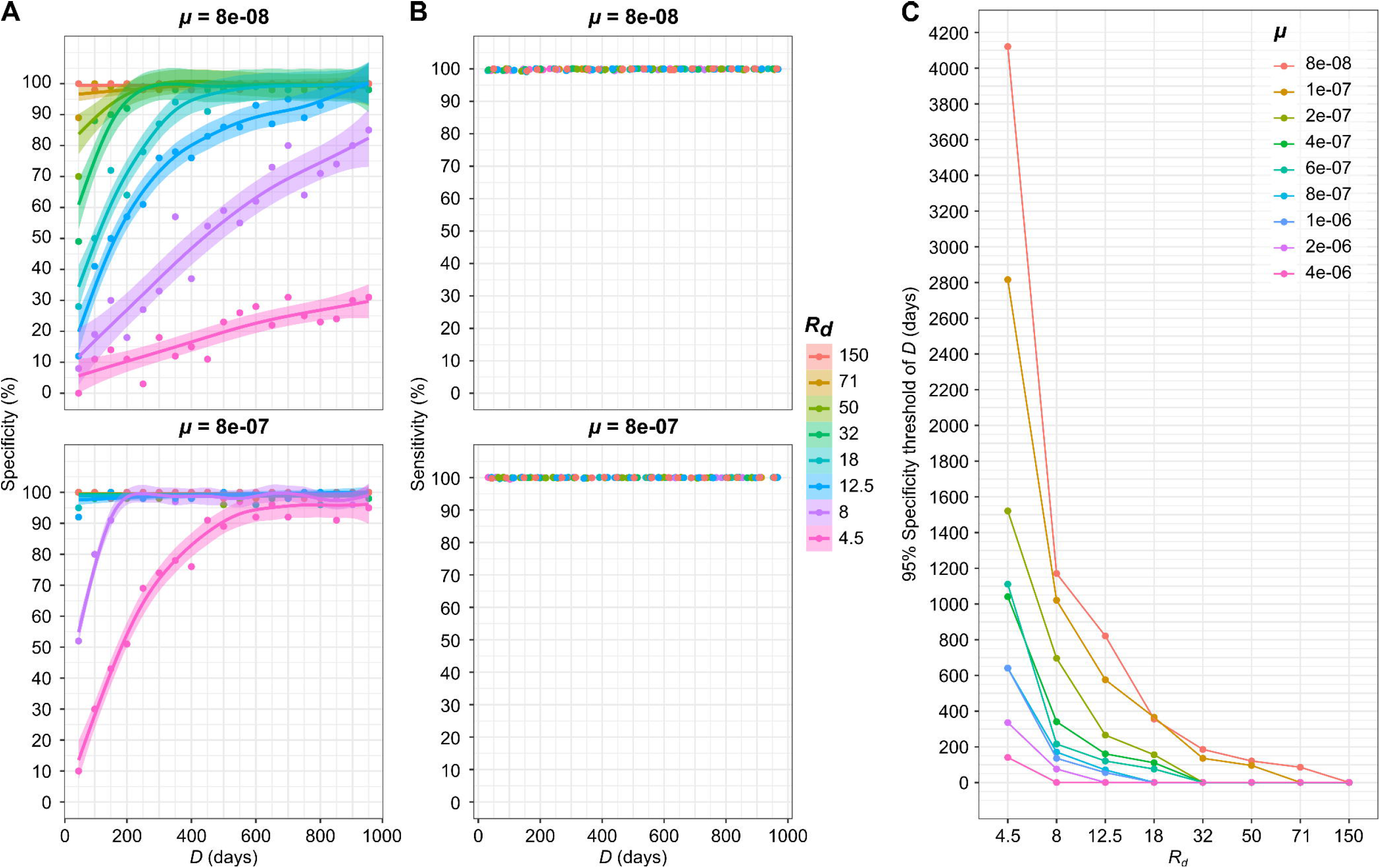
Assessment of the model’s ability to classify outbreak isolates from the simulation study. Specificity (**A**) and sensitivity (**B**) of isolates classification when *μ* = 8e-08 (top) or 8e-07 (bottom) substitutions per site per year. Each point provides specificity or sensitivity computed from 20 independent outbreaks simulated with the same input parameters, with *D* ranging from 50 to 1000 days (x-axis) and *R*_*d*_ (the ratio of evolution duration between non-outbreak and outbreak genomes) varying between 4.5 and 150 (colours). (**C**) 95% specificity threshold value of *D* as a function of *R*_*d*_ (x-axis), computed for 9 values of *μ* (colours).

We next evaluated whether the model and framework could accurately estimate the parameters *D* and *μ* from outbreaks data. To do so, we simulated synthetic outbreaks for which the *D* and *μ* values were known, and attempted to estimate one or the other. Regarding *D* estimation, all HPD include the true value, with higher values of *D* being associated with smaller 95% HPD (**Figure 3A**). Similarly, *μ* was adequately estimated, with best estimates being closer to the target value for higher *μ* values (**Figure 3B**). Because higher *D* and/or *μ* values lead in average to more SNPs, we indeed expected more precision in HPDs estimates in these cases.

**Figure 3.**
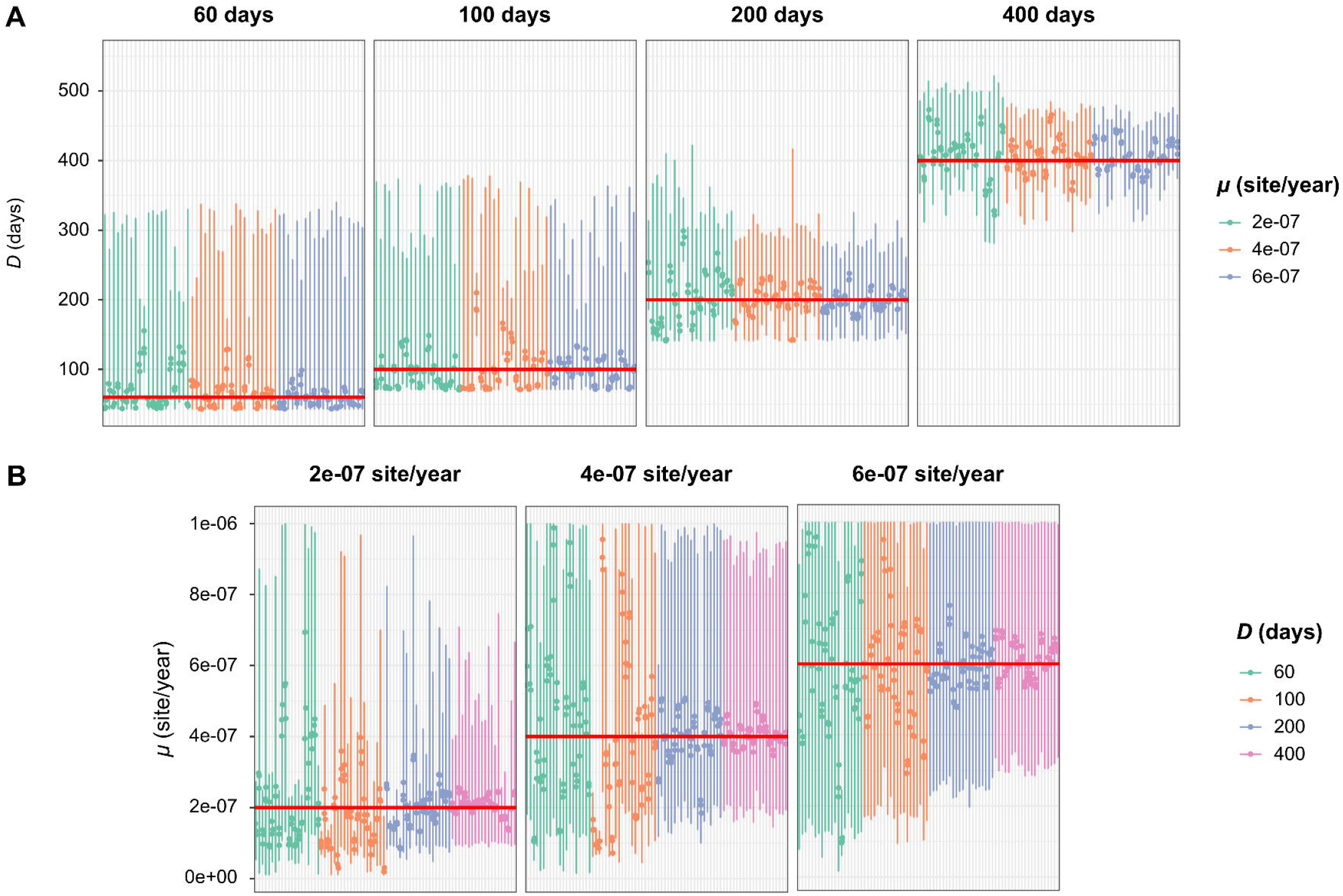
Assessment of the quality of the estimation of *D* and *μ* through simulation. Precision of the estimation of *D* (**A**) and *μ* (**B**) from simulated data generated using different values of *D* and *μ*. The sample size, defined as the number of observed samples and associated dates, was set to 0.2x*D*. On each panel, the upper banner and red line indicate the expected *D* (A) and *μ* (B) values used to generate the simulated data. For each of the 240 synthetic outbreaks analysed, three independent MCMC chains were run, and the three corresponding best estimates are shown (points). Vertical bars represent the average values of the minimum and maximum of the 95% credible interval of the 3 MCMC chains. Each colour corresponds to distinct values of *μ* or *D* used in the simulations (see keys).

We also investigated the impact of sampling density on estimation accuracy. Results suggest that poor sampling densities (*e*.*g*., 5%), when associated to low values of *D* and *μ* (therefore resulting in a low genetic diversity among samples), resulted in biased estimations of *D* and *μ*, which were generally overestimated (**Figure 4A** and **4B**). However, we show that sampling densities >10% led to unbiased estimations.

**Figure 4.**
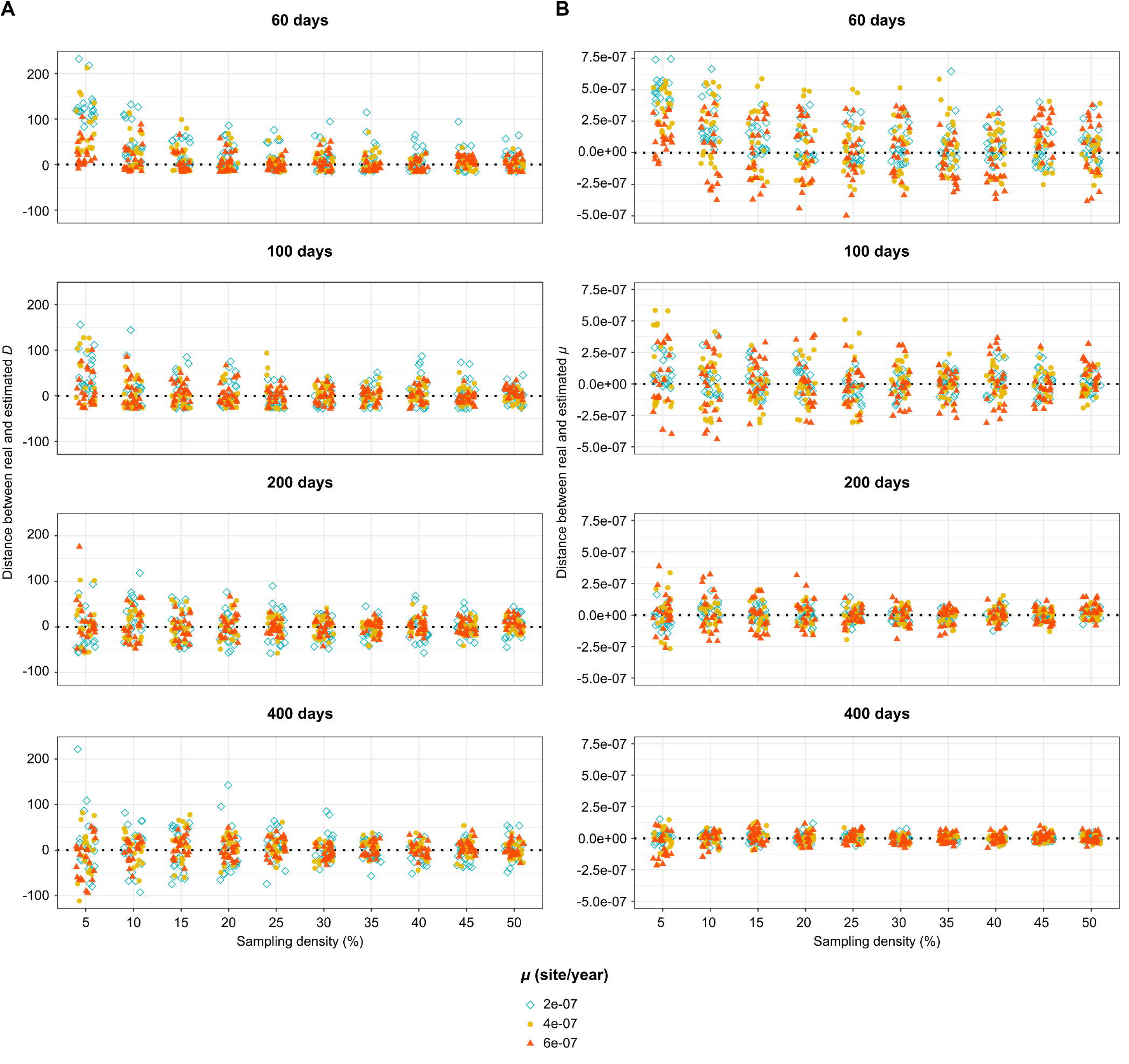
Impact of sampling density on the precision of the estimation of *D* and *μ*. The position of each symbol represents the difference between the expected and best estimates of *D* (A) and *μ* (B) for each of 2400 outbreaks simulated using combinations of 4 values of *D* (60, 100, 200 and 400 days; represented in rows) and 3 values of *μ* (2E-07: blue diamonds, 4E-07: orange circles, 6E-07, red triangles; values in substitutions per site per year). Sampling density represents the percentage of individuals sampled at each time step.

### Genetic threshold definition for published outbreak datasets

For each of the 16 published outbreaks, we applied our framework to estimate an expected outbreak-specific genetic threshold value (**Figure 5** provides the example of outbreak 11; see Supplementary appendix figures S1 to S16 for all outbreaks). We found that, for 14 out of 16 outbreaks, the classification of isolates as being outbreak-related or sporadic is consistent with previously reported results. Four of these outbreaks included outliers (outbreaks 1, 4, 12 and 16), which are correctly classified beyond the threshold of exclusion by our model, except for one isolate of outbreak 4 (**Table 1; Fig S4;** note that outbreak 4 comprised three different co-contaminating genetic clusters [20]; here the defined outbreak strain was ST528). Ten other outbreaks (2, 3, 5, 6, 7, 9, 10, 13, 14 and 15) have no sporadic cases, and our framework correctly clusters all suspected isolates as outbreak-related.

**Figure 5.**
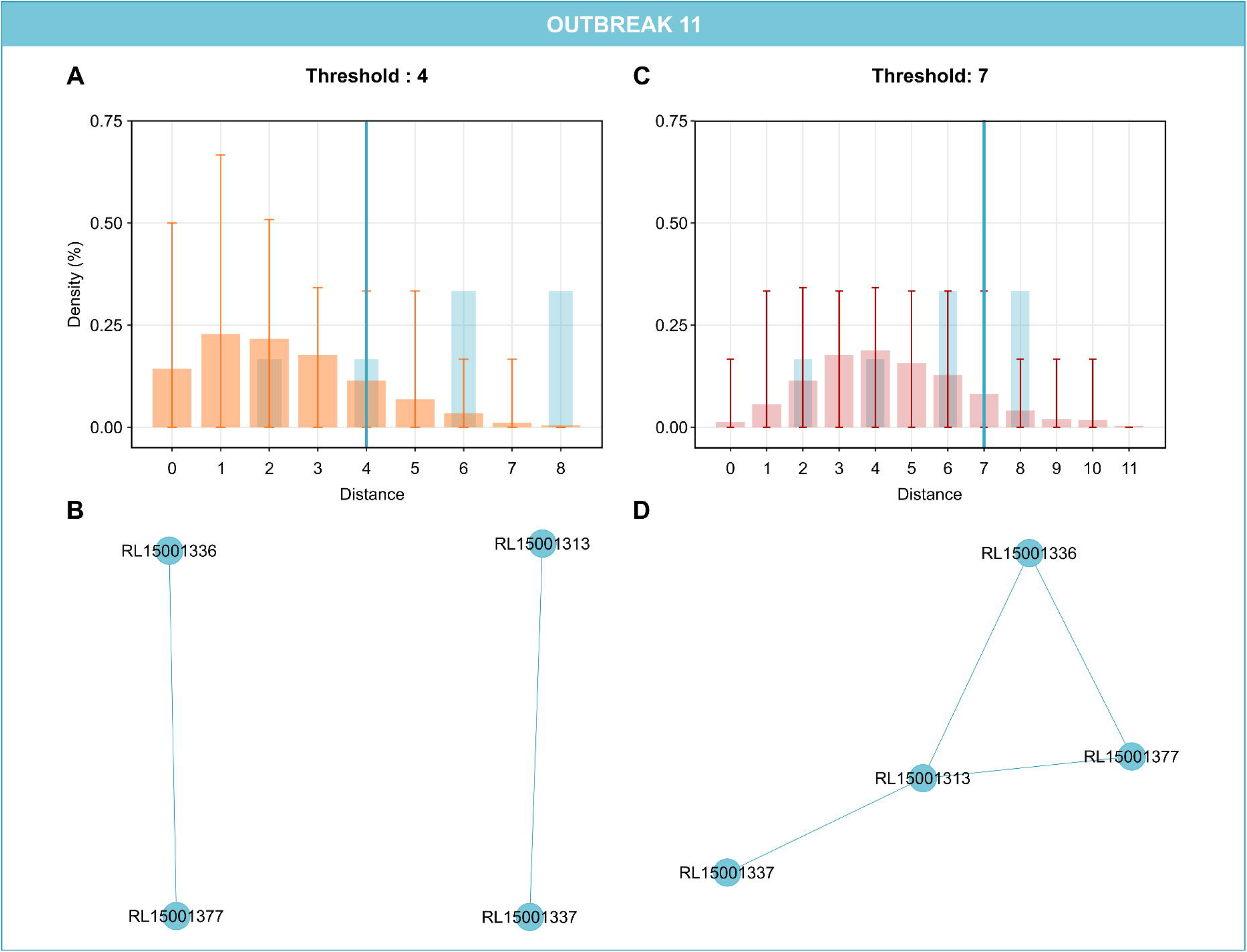
Distance threshold derived from the modelling framework, and its effect on clustering: example of outbreak 11. Panels A and C show the cgMLST distance distributions: observed distribution (blue, panels A and C), simulated distribution without estimation (Orange, panel A), and simulated distribution using the estimated duration of outbreak (red, panel C). Error bars represent the interval of prediction at 95% of 100 simulations. Blue vertical lines correspond to the derived distance threshold defined here as the 99^th^ percentile of the distributions (A: from the observed distribution; C: from the simulated distribution using the estimated duration of outbreak). Panels B and D show the single-linkage clusters resulting from the derived distance threshold corresponding to panels A and C, respectively.

For two of the 16 outbreaks, our model leads to different conclusions compared with previous results. In outbreak 8 (*L. monocytogenes*, beef), two isolates are classified as outliers by our model, whereas they were initially classified as outbreak-related [24]. In outbreak 11 (*L. monocytogenes*, ox tongue), two isolates came from food and two others from humans. Our algorithm separates food samples in one cluster and human samples in another cluster, whereas the isolates were initially grouped based on epidemiological and genetic evidence: here, the threshold inferred by our model was smaller.

When evaluating the influence of outliers on the inferred threshold by removing them from the analysis we find that, in all cases, the outliers do not affect the outbreak threshold. For outbreak 1, 4 and 16, this removal does not change the threshold value but improves the fit between the pairwise SNP distance distribution of the data and the simulated one (**Supplementary Table S2**).

### Estimation of *D* and *μ* values from real outbreaks, and impact on outbreak definition

For each of the 16 above outbreaks, we used our framework to estimate outbreak duration *D* and substitution rate *μ* (called *D*_*estimated*_ and *μ*_*estimated*_) separately, and used these values (instead of *D*_*lit*_ and *μ*_*lit*_ used above) for the inference of the genetic distance threshold. Results are provided in **Table 1**.

For 11 of the 16 outbreaks, the estimated HPD intervals include *D*_*lit*_. For the 5 remaining, we find higher *D*_*estimated*_ values compared with previously reported *D*_*lit*_ (**Figure S6** and **Table 1**). Regarding *μ*, for nine outbreaks HPD intervals include their corresponding *μ*_*lit*_, whereas for only one outbreak *μ*_*estimated*_ is lower than *μ*_lit_ (outbreak 2) and the six remaining outbreaks lead to a higher estimated *μ*_*estimated*_ compared with *μ*_*lit*_. It is important to note that the *D*_*estimated*_ 95% HPD is also higher than *D*_*lit*_ for these same 6 outbreaks (**Table 1**).

After reanalysing the outbreaks using our *D*_*estimated*_ and *μ*_*estimated*_ values, we observe that the newly obtained thresholds do not affect the attribution of isolates to the outbreak or sporadic categories in most cases, with three exceptions. First, for outbreak 4, using *D*_*estimated*_ or *μ*_*estimated*_ increases the threshold from 4 to 11 SNPs, leading to add the previously missing isolate but still excluding the outliers. Second, for outbreak 15, a decreased genetic threshold (4 SNPs instead of 5, in both independent estimations analyses for *D*_*estimated*_ and *μ*_*estimated*_) leads to the exclusion of one isolate. Third, for outbreak 11, the genetic threshold is increased from 4 SNPs to 7 and 10 SNPs (using *D*_*estimated*_ and *μ*_*estimated*_ respectively), leading to group all isolates from food and human samples (**Figure 5**). We also observe that in most cases, using the estimated values of *D* and *μ* improves the fit of the genetic distance distribution, with two exceptions (**Table S2** in the supplementary appendix).

## Discussion

Molecular surveillance contributes to identify common exposure to a specific source even when dates and places of infections are distant [25–27]. Given the large differences existing among outbreaks, it is being increasingly recognised that no single-species threshold can be applied to distinguish between outbreak and non-outbreak isolates. To our knowledge, Octavia and colleagues (2015) were the first to attempt to model the expected genetic distance among food outbreak isolates. Although the authors incorporated mutation rate and outbreak duration in their model, they did not use the actual sampling dates. Consequently, their proposed thresholds depend on strong assumptions as to the actual duration of the outbreak (referred to as the *ex-vivo/in-vivo* evolution time by these authors). Stimson *et al*. [14] modelled the number of transmissions that separates infection cases, using a probabilistic model that incorporates the transmission process in addition to mutation rate and timing of infections. Because it models between-host transmission, this approach does not apply to point-source food outbreaks. Lastly, Coll *et al*. [28] aimed at defining a SNP threshold above which transmission of *S. aureus* between humans can be ruled out, by incorporating the timing of transmission and within-host diversity. This evolutionary modelling approach provides a robust SNP cut-off applicable to this specific ecological situation.

We propose an original evolutionary approach to the ‘single strain’ threshold conundrum by incorporating epidemiological and microbiological specifics of each outbreak. Our model is supported by a high sensitivity (>90%) of isolates classification and by the results of analyses of 16 real-life published datasets from foodborne outbreaks, which led to consistent results in most cases and enabled to refine outbreak analysis in two cases.

The simulation study showed that our model performed well at grouping outbreak cases. We also observed that as *D* and *μ* increased, the estimated genetic threshold was more accurate: the model specificity increased with genetic diversity. This is akin to higher resolution typing methods being better at discriminating related and non-related cases. We also found an impact of the evolutionary distance between outbreak and sporadic isolates on model specificity, consistent with the known uncertainty in ruling out sporadic cases for genetically homogeneous pathogens. In addition, we found that the sampling density is important, as it influences the number of observed genetic differences: outbreaks with low diversity will require more samples to capture enough pairwise differences for estimation purposes.

Our model assumes a constant population, to avoid increasing execution time with growing bacterial populations. Because the population *N* remains constant over time, this number must be chosen high enough to capture all the diversity through our sampling process. Indeed, we simulated the sampling processes and did not analyse the whole *N* population. Because λ, the Poisson parameter, is defined as a function of *N*, a number of 500 or 1000 is usually enough to capture all bacterial diversity, but higher values should be tested further when extreme substitution rates or duration are explored.

In most outbreak investigations, the time since source contamination is unknown, and the underestimation of *D* is a common risk given the possibility of cryptic transmission and unreported cases having occurred prior to actual outbreak detection [29]. Prior knowledge of *μ* is also subjected to uncertainty: this parameter strongly depends on the species but also on the strain [30] and on other conditions (*e*.*g*. temperature, cellular stress). We showed that, although the estimates were largely consistent with epidemiological information, estimated *D* and *μ* were often larger. As *D* and *μ* both affect the expected genetic diversity in the same direction, it is impossible to decide whether it is the rate, or the duration, that was higher than initially suspected. We suggest that, in the absence of evidence for higher *μ*, fixing it and estimating *D* may provide important clues regarding prior cryptic transmission. Considering higher *D* values than suggested by case recognition is clearly relevant for epidemiological investigations of outbreaks, as it widens the considered time window and may lead to identify initially unsuspected sources of contamination.

The analysis of the 16 published outbreaks led to the definition of genetic thresholds that were largely consistent with epidemiological evidence. For outbreaks 4 and 11, groupings were discordant, as a lower threshold than initially used was inferred by our model. However, when estimating the duration or substitution rate with our framework, higher values were observed for both outbreaks, thus leading to group samples consistently with epidemiological evidence. Outbreak 11 involved foodborne listeriosis with contaminated food where the two food samples differed by 9 SNPs from the human samples, themselves separated by 2 SNPs. The two food samples were isolated from two food outlets that had the same meat producer. Because the incubation period of listeriosis is between 3 and 70 days and because intermittent *L. monocytogenes* contamination during the production was observed [31], the duration of contamination *D* might have been higher than initially defined, suggesting that the true common ancestor of food and human isolates was in fact older than initially estimated from the original publication. This illustrates the value of our estimation framework to inform epidemiological investigations. Interestingly, when using model-estimated duration of outbreak or substitution rate, we often observed an improved fit of the pairwise distance distributions (**Table S2**).

For outbreak 8, low sequence data quality was observed for three genomes [24], including the two genomes excluded from the outbreak by our model. Low quality data may have artificially inflated their genetic distinctness, which underlines the importance of input sequence data quality.

It is important to highlight the following limitations. First, all presented results were generated by initialising the models with a fully homogeneous ancestral population. However, the contaminating population may be slightly heterogeneous if it has a non-negligible population size and had itself already evolved previously. In these cases, *D* might be interpreted as incorporating the diversification time before source contamination. Second, we only modelled mutation, neglecting other evolutionary processes such as recombination. Detection of recombination among very closely related isolates is very challenging and its impact would be limited. However, recombination with genetically distinct co-contaminants might occur and recombined chromosomal regions should be removed from the analysis, especially when using SNP-based analyses (by design, MLST moderates the impact of homologous recombination). Third, the model does not incorporate demographic events within the contaminated source, including population bottlenecks, which are potentially common in food processing chains but which would be challenging to infer and to model. Finally, the framework is designed for a single evolving population derived from a single bacterial ancestor. When there is more than one contaminating genotype, our framework could be used separately for each of these.

### Conclusions

We describe an innovative approach to the ‘single strain’ definition using pathogen genomic data by considering the most relevant features of specific outbreaks to define a credible genetic distance threshold. This definition is grounded in evolutionary biology and alleviates the need for *a priori* defined thresholds, which are not justified theoretically and may be inappropriate in most cases. The inferred outbreak-tailored genetic thresholds provide a reliable, non-arbitrary way to define epidemiologically related infection cases and to exclude non-related sporadic strains. This approach is fast and easy to use. The additional ability to estimate outbreak duration should also prove useful for point-source disease outbreak studies, by providing a credible temporal window for epidemiological investigations aiming at identifying and eliminating the sources of contaminations.

## Acknowledgements

We acknowledge the financial support of the MedVetKlebs project, a component of European Joint Programme One Health EP, which has received funding from the European Union Horizon 2020 Research and Innovation Programme (Grant N. 773830). We thank Xavier Didelot and Olivier Tenaillon for critical feedback on an earlier version of the manuscript, and Chiara Crestani for help in manuscript and figures formatting.

## Authors contributions

S.B. conceived the presented approach. A.D., L.O. and S.B. designed the model and the computational framework. A.D. programmed the model and analysed data and simulations. A.D., L.O. and S.B. interpreted the results. A.D., L.O. and S.B. wrote the manuscript. The three authors read and approved the final manuscript.

