## Supplementary Appendix for "Is it the same strain? Defining genomic epidemiology thresholds tailored to individual outbreaks"

**Contents**

Supplementary text	pages 2-6

Supplementary tables	page 7

Supplementary figures	pages 9-26

Supplementary references	page 27

**Supplementary text**

**Methods**

**Evolutionary model and definition of the outbreak genetic distance threshold**

Our evolutionary formalization is based on a Wright-Fisher forward model of haploid infectious agent evolution using a constant population. Here, we set the population size N = 500 but any large-enough number from which samples will be drawn randomly would be suitable. Within our modelling framework, the five distinct properties characterising the infectious agent responsible for an outbreak are: i) *L*, the genome length (in base pairs, bp) or the average length of genes if gene-by-gene genotyping is used, such as in multilocus sequence typing (MLST) approaches; ii) *g*, the number of genes*; g* is equal to 1 when genome length is considered*.* For core-genome SNP data, *L* should be set to the length of the core genome considered, rather than the whole genome length; iii) *μ*, the number of substitutions per site per year; iv) *D,* the duration (in days) of the outbreak. *D* corresponds to the duration (in days) of (unfrozen) source contamination during which evolution and exposure of cases takes place, until the last recorded sample; and v) *S_d_*, the set of sampling dates of isolates. Although *D* could potentially be approximated directly from *S_d_* in some cases, it is useful to define *D* as being larger than the largest time interval within *S_d_*, for example when suspicion exists that the source was contaminated much earlier than the first available sample. We define the end of the simulation process as the last sampling date, whilst the other sampling dates are defined backwards in time relative to their time distance to the last sample.

The simulation is run during D days using one-day time steps, drawing the number of substitutions for the whole bacterial population at each time step. The evolutionary process is repeated for all time steps until D is reached. We assumed that the number of substitutions introduced at each time step during our simulation follows a Poisson distribution with parameter $\lambda$ computed as:

*µ*~Poisson (*λ*)

$$\lambda= \frac{\mu}{365} NLg$$

(1)

where *N* is the population size, *L* is the length of the sequence (or the gene in MLST contexts) and *g* the number of genes. A given individual can receive one or multiple substitutions at a given time step. All substitutions are considered unique (*i.e.,* lead to a new allele). At the end of this process, *n* individuals are randomly sampled to match the size of the studied dataset at sampling dates corresponding to those in the observed sample set *Sd*. A distribution of pairwise genetic distances is generated on these sampled individuals. We then chose the genetic threshold value as an upper percentile, *e.g*., here the 99^th^ percentile, of this distribution. If the genetic distance of an isolate to at least one other member of the group is lower or equal to the threshold value, this isolate is considered as a member of the outbreak group. Otherwise, the isolate is considered an outlier.

**Estimation of the model’s ability to discriminate between outbreak and outlier samples using simulated outbreaks**

We applied our framework to a series of 171 simulated outbreaks generated with 19 different values of *D* each combined with 9 values of *μ* and including simulated sporadic isolates. Synthetic datasets were built in two steps by generating on the one hand, a series of samples corresponding to a simulated outbreak; and on the other hand, a series of samples corresponding to simulated outliers, here called sporadic isolates. These latter were drawn from another simulated outbreak defined by *D_sporadic_*; this approach enables to challenge the model with varying degrees of distinctness between outbreak and sporadic isolates, with the latter resulting from distinct durations of evolution since a common ancestor shared with the outbreak isolates. Details can be found in **Table S1**.

For a given analysis, we defined true positives (TP) defined as the real outbreak cases classified as a part of the outbreak, true negatives (TN) as sporadic cases classified as being excluded from the outbreak, false negatives (FN) as real outbreak cases excluded from the outbreak by the model, and false positives (FP) as the sporadic isolates misclassified as members of the outbreak. We then computed sensitivity (*Se*) and specificity (*Sp*) of our approach based on the following formulae: $Se= \frac{TP}{TP+FN}$ and *Sp* $= \frac{TN}{FP+TN}$.

We set *g* = 1, *L* = 4,800,000 bp and *N* = 500. For each *D*, *μ* and *D_sporadic_* values combination, 20 independent outbreaks with 5 sporadic cases each were stochastically generated. For each outbreak, samples dates *S_d_* were randomly selected such as *S_d_* ~ *Uniform*(0, *D*) and *S_d_* ~*Uniform*(*D_sporadic_*-*D*, *D_sporadic_*).

We defined *R_d_* as the ratio of evolution duration of the sporadic population to the evolution duration of the outbreak population (*i.e.*, $R_{d}= \frac{D_{sporadic}}{D}$). To assess to which extent the genetic distance between the outbreak and sporadic isolates can alter the quality of our results, we investigated the impact of varying the ratio of evolution duration *R_d_*. Eight *R_d_* values were explored: 4.5, 8, 12.5, 18, 32, 50, 71 and 150. For each *R_d_*, 20 outbreak simulations were run, and this for all 171 distinct combinations of *D* and *μ*. In total, 3420 outbreaks and 27,360 outbreak-sporadic synthetic datasets were simulated.

For each *R_d_* and *μ* combination values, independent Generalized Additive Model (GAM) regression were applied to investigate the link between specificity and duration. To assess the minimum *D* required to guarantee 95% specificity, we collected the *D*-value corresponding to the 95% specificity threshold from the prediction of the GAM regression.

**Estimation of the duration and substitution** **rate of outbreaks**

To address uncertainty on the time since source contamination and evolutionary rate, we embed our model into a statistical framework to estimate either the duration of the outbreak (*D*) or its substitution rate (*μ*) (**Figure 1B**). We estimate *D* and *µ* from the observed pairwise genetic distance matrix by minimizing the distance described below, using a Markov Chain Monte Carlo (MCMC) with the Delayed Rejection Adaptive Metropolis (DRAM) algorithm from R package FME [19]. For each possible set of parameters (*L*, *g*, *S_d_*, *D* or/and *μ*), the least squares distance (LSD) is used to evaluate the match between the observed pairwise genetic distance matrix and the expected one associated to the parameters set. For a given *L*, *g*, *S_d_*, *D* or/and *μ*, the expected distribution of pairwise genetic distances is obtained by simulating population evolution using the previously presented model. Estimated parameters are provided by computing the mean of the best posterior estimates obtained for 3 independent MCMC chains with the average 95% highest posterior density (HPD).

From the simulated distribution of pairwise genetic distances (for a given *L*, *g*, *S_d_*, *D* or/and *μ)*, the least squares distance is defined as (under the 99^th^ percentile, as the upper values are considered as non-outbreak related):

$$LSD= \sum_{i=1}^{n} (y_{i}-{y'}_{i})^{2}$$

(2)

Where *n* is the highest difference in term of number of SNP or cgMLST under the 99^th^ percentile between observed data *y* and expected data from simulations *y’*.

Because of the simulation stochasticity, for each MCMC iteration, 20 independent model simulations are run and averaged to build the expected distribution. In total, 10,000 iterations of the MCMC are run, with the 2000 first ones being discarded as a burn-in. All MCMC chains are run with the modMCMC function from the FME package version 3.5.3 of R [19].

*Validation*. To assess the ability of the model to properly estimate *D* and *μ*, we run our framework on simulated outbreaks for which parameters were known. *L*, *N* and *g* were fixed as above. Outbreaks generated using combinations of (D, *μ*) were assessed, by varying durations over *D* = 60, 100, 200, 400 days and substitution rates over *μ =* 2E-07, 4E-07 and 6E-07 substitutions per site per year. Because the number of sampled genomes could affect estimation quality, and to assess to which extent this was the case, the number of samples was also varied in the analysis. For each outbreak, different sampling densities were evaluated, the number of sampled days ranging from 5 to 50% of *D*. For each combination of parameters, 20 individual outbreaks were run, leading to 2400 different simulated datasets. For each outbreak, we estimated *D* and *μ* separately with three independent MCMC chains.

**Descriptions of published outbreaks investigated with the modelling framework**

Outbreak 1 (1) was caused in November 2016 by *Salmonella enterica* serovar Typhimurium phage type DT170. The possible food source of contamination was a chocolate mousse. Among 47 cases, 13 isolates were sequenced from patients. One isolate differed by 12 SNPs and another by five SNPs, which was interpreted as at least three distinct strains being present in the food source. We used the authors’ high substitution rate and high duration of outbreak for *D_lit_* and *μ_lit_*.

Outbreaks 2 and 3 also involved *Salmonella* Typhimurium in Australia (2). These two outbreaks occurred between January and May 2014 in metropolitan Sydney. Outbreak 2 was related to a chicken liver pâté and outbreak 3 to a hot bread shop. At least 25 isolates for outbreak 2 and 20 isolates for outbreak 3 were sequenced and available (as well as some sporadic cases that we did not consider in the present paper). We used the same substitution rate for *μ_lit_* as for Outbreak 1 and duration of outbreaks *D_lit_* were chosen as the duration between the first and the last sample dates (+24H).

Outbreaks 4 (3) and 5 (4) involved *Campylobacter jejuni* infections caused by contaminated chicken liver pâté in Australia and milk in Finland, respectively. Outbreak 4 occurred from the end of October to the beginning of November 2013; 9 isolates (7 from humans and 2 from food) were sequenced. Three distinct clusters were identified, ST528 (five isolates), ST535 (two isolates) and ST991 (two food isolates); here we focused on ST528 isolates. Outbreak 5 involved two *C. jejuni* isolates from a milk tank, two from humans and two from dairy cows, collected from December 2002 to January 2003. Substitution rates from Wilson et al. (5)were used for outbreaks 4 and 5 as *μ_lit_*. We used the time difference between the first day of paté’s preparation and the last swab day (+24H) for *D_lit_* in outbreak 4. We used *D_lit_* as the duration between the first and the last sample dates (+24H) for outbreak 5.

Outbreaks 6 (6) and 7 (7) were caused by *Escherichia coli* O104:H4 and O157, respectively. Outbreak 6 involved cases in Germany and France between May to July 2011 and caused bloody diarrhoea and haemolytic uremic syndromes. This outbreak was epidemiologically linked to contaminated sprouts. Four isolates were sequenced from German cases and 11 from French cases. In outbreak 7, *E. coli* O157 was linked to unwashed vegetables in the UK in 2011. A total of 10 isolates were available. Substitution rates from Grad et al. (8) and Reeves et al. (9) were chosen for *μ_lit_* respectively. For outbreak 6, the duration of outbreak *D_lit_* was taken as the difference between first-55 days and last sample date because a traveller was suspected to be the cause of an introduction. For outbreak 7, we used the duration between the first and the last sample dates (+24H) for *D_lit_*.

Finally, nine distinct outbreaks, labelled by us 8 to 16 (**Table 1**), were caused by *Listeria monocytogenes* (10). These outbreaks were linked to beef, crabmeat, sandwiches, ox tongue, an unknown source, rakfisk, foie gras, cheese and brie cheese, respectively, from 2011 to 2014 in four unknown countries referenced as B, T, X and C. Among the 10 isolates in outbreak 12, one was attributed to a separate clonal complex by MLST and another one (from milk) had no epidemiological evidence of being linked. In outbreak 16 from Brie cheese, the 25 isolates were a mix of outbreak and background isolates, with only 11 isolates ultimately being attributed to the outbreak. For these 9 outbreaks, the substitution rate from Halbedel *et al*. (11) was used as *μ_lit_*. For *D_lit_*, the difference between first and last sample dates (+24H) was used for outbreak 8, 9, 10, 11, 12 and 14 and we added 6 months for outbreak 13 (time for fish ripening), 4 and 1 months for outbreak 15 and 16 (time for cheese ripening) and 2 months for outbreak 11.

**Computational demands of our modelling framework**

Execution time is very dependent of the parameters. We assessed the execution on a computer with 32Go of RAM and a processor intel® cor™ i7-10610U CPU@ for 4 outbreaks; (i) outbreak 1 with classical *D* and *μ*, (ii) outbreak 6 with a shorter *D* and a larger μ, (iii) outbreak 10 with longer *D* and lower *μ* and (iv) outbreak 5 with a shorter *D* and the highest *μ*. Estimating the genetic threshold (with the classical 100 simulations) is very fast and takes around 0.1, 0.1, 0.2 and 1.6 seconds for outbreaks 1, 6, 10 and 5 respectively. Nevertheless, estimation of *D* or *μ* with one MCMC chain is much longer and takes around 40, 47, 100 and 1500 seconds for the estimation of *μ* as well as 46, 70, 43, and 300 seconds for the estimation of *D* for outbreaks 1, 6, 10 and 5 respectively. The more *D* or *μ* were high the more the estimation was long. Especially, *μ* had a more important impact on the execution time.

**Supplementary tables**

**Table S1. Simulations used to validate the model.**

|  | **Number of *D* values used** | **Number of *µ* values used** | **Number of *R_d_* values used** | **Number of sampling density values** | **Number of repetitions (per value set)** | **Total number of simulated outbreaks** |
| --- | --- | --- | --- | --- | --- | --- |
| **Simulated outbreaks for threshold definition** | 19 (from 50 to 950 days) | 9 (from 8e-08 to 4e-06 substitutions per site per year) | 8 | 1 (20 outbreak samples and 5 outliers) | 20 | 27 360 |
| **Simulated outbreak for estimation of *µ* and *D*** | 4 (from 60 to 400 days) | 3 (from 2e-07 to 6e-07) | 0 | 10 (from 5% to 50%) | 20 | 2 400 |

**Table S2. Kolmogorov-Smirnoff (noted *D_KS_*) test statistic for each of the 16 published outbreaks**.

|  | ***D_KS_*** | | |
| --- | --- | --- | --- |
| **Outbreak** | ***D_lit_* and *μ_lit_*** | ***D_estimated_*** | ***μ_estimated_*** |
| 1 | 0.33 | 0.25 | 0.25 |
| 2 | 0.5 | 0.5 | 0.5 |
| 3 | 0.33 | 0.33 | 0.33 |
| 4 | 0.8 | 0.58 | 0.58 |
| 5 | 0.41 | 0.50 | 0.47 |
| 6 | 0.5 | 0.21 | 0.14 |
| 7 | 0.33 | 0.33 | 0.33 |
| 8 | 0.33 | 0.33 | 0.75 |
| 9 | 0.38 | 0.36 | 0.33 |
| 10 | 0.33 | 0.4 | 0.33 |
| 11 | 0.6 | 0.62 | 0.64 |
| 12 | 0.57 | 0.57 | 0.4 |
| 13 | 0.5 | 0.5 | 1 |
| 14 | 0.67 | 0.17 | 0.33 |
| 15 | 0.33 | 0.2 | 0.4 |
| 16 | 1 | 0.75 | 0.6 |

Outbreak numbers refer to Table 1. We analyzed the fit of the distribution of genetic distances obtained when using either *D_lit_* and *μ_lit,_* or *D_estimated_* or *μ_estimated_*, using the Kolmogorov-Smirnoff test statistic (*D_KS_*). *D_KS_* varies from 0 to 1. Small *D_KS_* lead to a better fit. The use of MCMC-estimated values instead of those defined from the literature led to a better fit for outbreaks 4, 6 and 16 **(see also Supplementary Figures S4, S6, S16).** For outbreaks 14 and 15, only *D_estimated_* improved the fit compared to both *D_lit_* and *μ_lit_* (**Supplementary Figure S10, S14 and S15**). For outbreak 12, a better fit was observed with *μ_estimated_* compared to *D_lit_* and *μ_lit_* (**Supplementary Figure S12**). However, the use of *μ_estimated_* lead to a less good fit for outbreaks 8 and 13 (**Supplementary Figure S8 and S13**). Indeed, the estimation of μ lead to the addition of one isolate in outbreak 8 while it did not change the threshold of outbreak 13. The estimation of *μ_estimated_* lead to increase the threshold from 2 SNPs to 7 SNPs but the fit of the genetic distance distribution was worst. Finally, no significant improvement or decline of the fit was observed for outbreaks 1, 2, 3, 5, 7, 9 and 11.

**Supplementary figures**

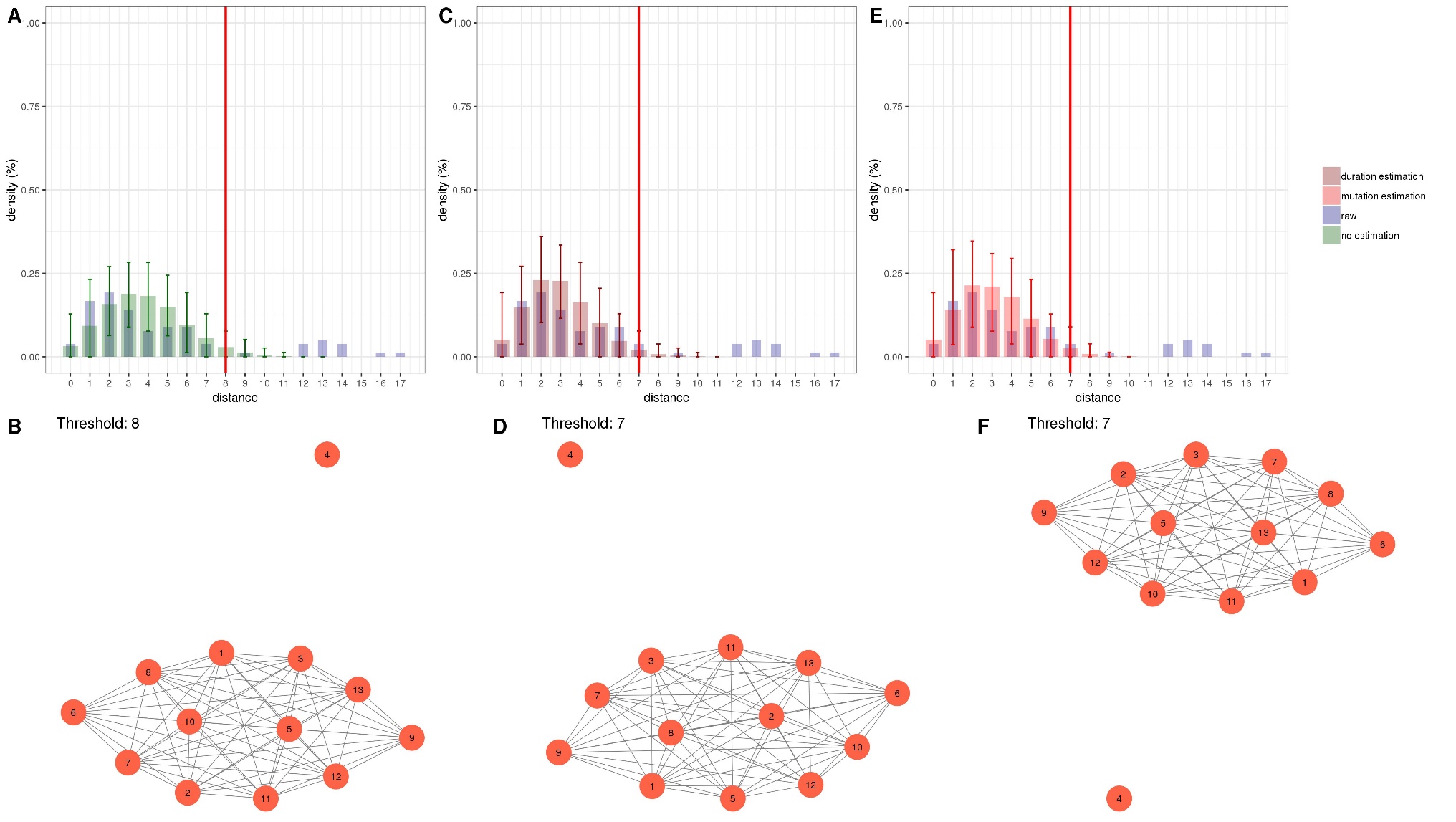

**Figure S1**. **Analysis of outbreak 1: Comparison between distance thresholds derived from the original publication on the outbreak, and from the current modelling framework.**

In A, C and E are presented the SNP distance distributions: observed (raw) distribution (blue), simulated distribution without estimation (green), simulated with the estimated duration of outbreak (dark red) and simulated with the estimated evolutionary rate (substitution, red). Error bars represent the interval of prediction at 95% of 100 simulations. Red vertical lines correspond to the derived distance threshold. In B, D and F are presented the resulting single-linkage clusters according to the derived distance threshold, defined here as the 99^th^ percentile of the simulated distributions with values corresponding to panels A, C and E, respectively.

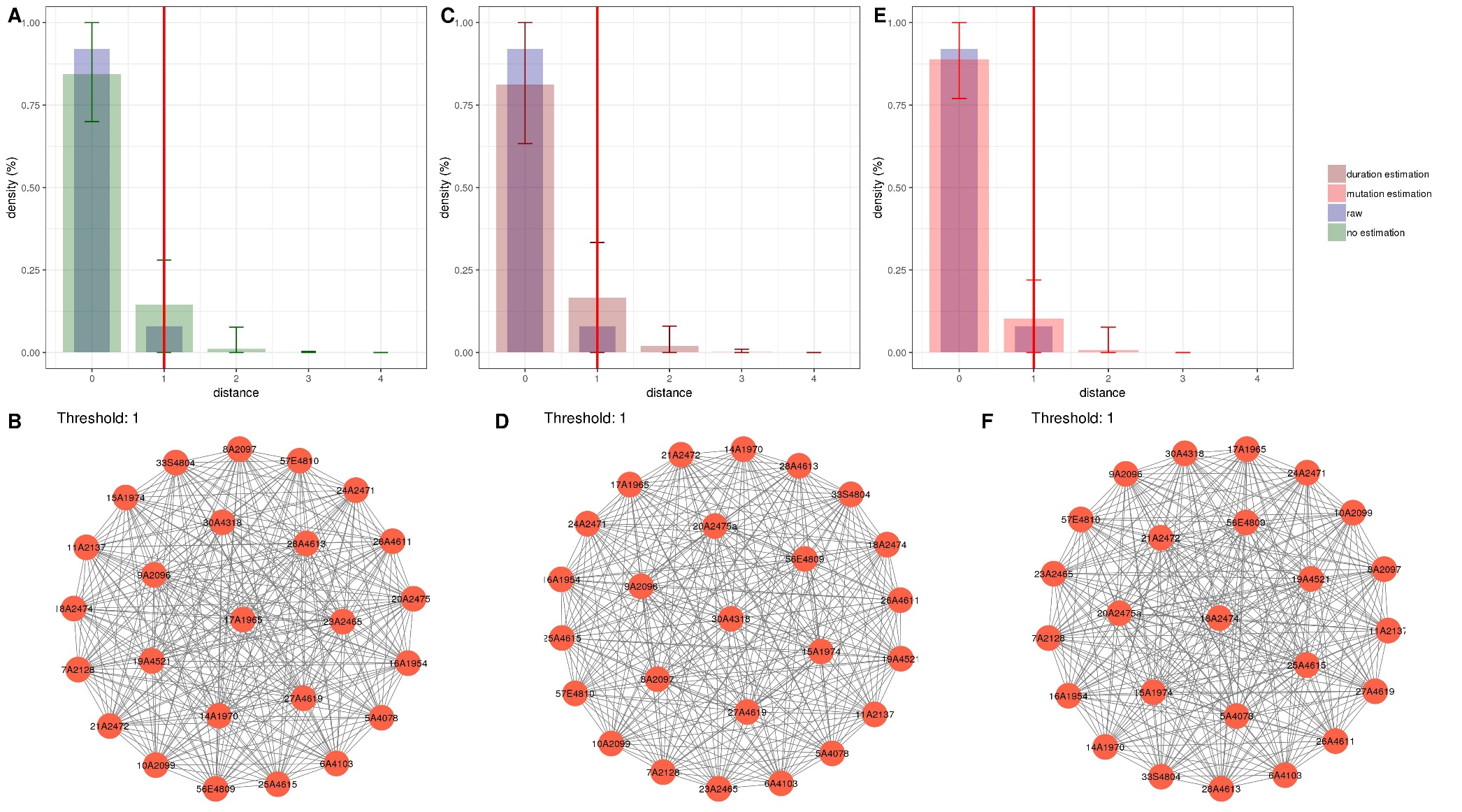

**Figure S2. Analysis of outbreak 2: Comparison between distance thresholds derived from the original publication on the outbreak, and from the current modelling framework.**

In A, C and E are presented the SNP distance distributions: observed (raw) distribution (blue), simulated distribution without estimation (green), simulated with the estimated duration of outbreak (dark red) and simulated with the estimated evolutionary rate (substitution, red). Error bars represent the interval of prediction at 95% of 100 simulations. Red vertical lines correspond to the derived distance threshold. In B, D and F are presented the resulting single-linkage clusters according to the derived distance threshold, defined here as the 99^th^ percentile of the simulated distributions with values corresponding to panels A, C and E, respectively

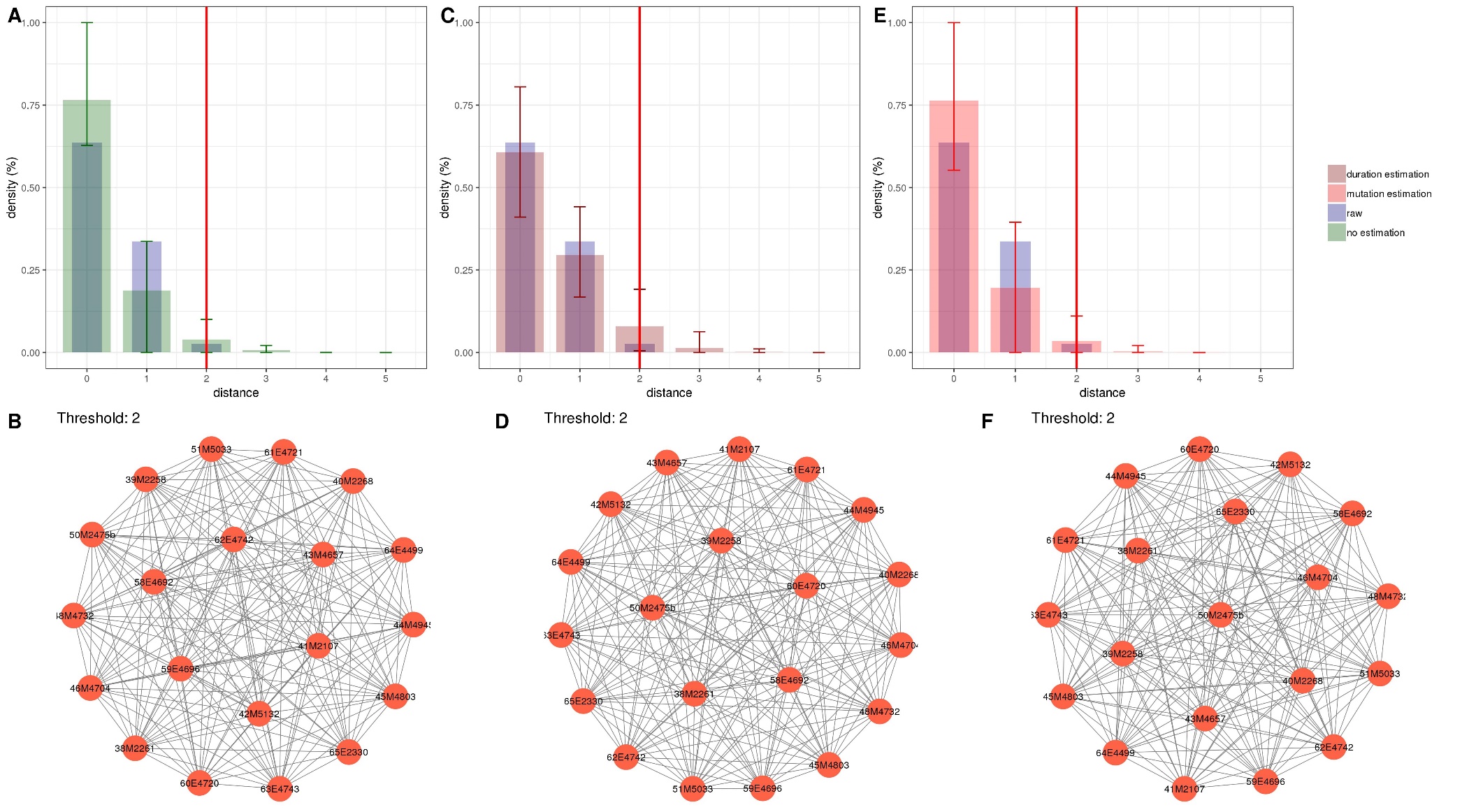

**Figure S3. Analysis of outbreak 3: Comparison between distance thresholds derived from the original publication on the outbreak, and from the current modelling framework.**

In A, C and E are presented the SNP distance distributions: observed (raw) distribution (blue), simulated distribution without estimation (green), simulated with the estimated duration of outbreak (dark red) and simulated with the estimated evolutionary rate (substitution, red). Error bars represent the interval of prediction at 95% of 100 simulations. Red vertical lines correspond to the derived distance threshold. In B, D and F are presented the resulting single-linkage clusters according to the derived distance threshold, defined here as the 99^th^ percentile of the simulated distributions with values corresponding to panels A, C and E, respectively

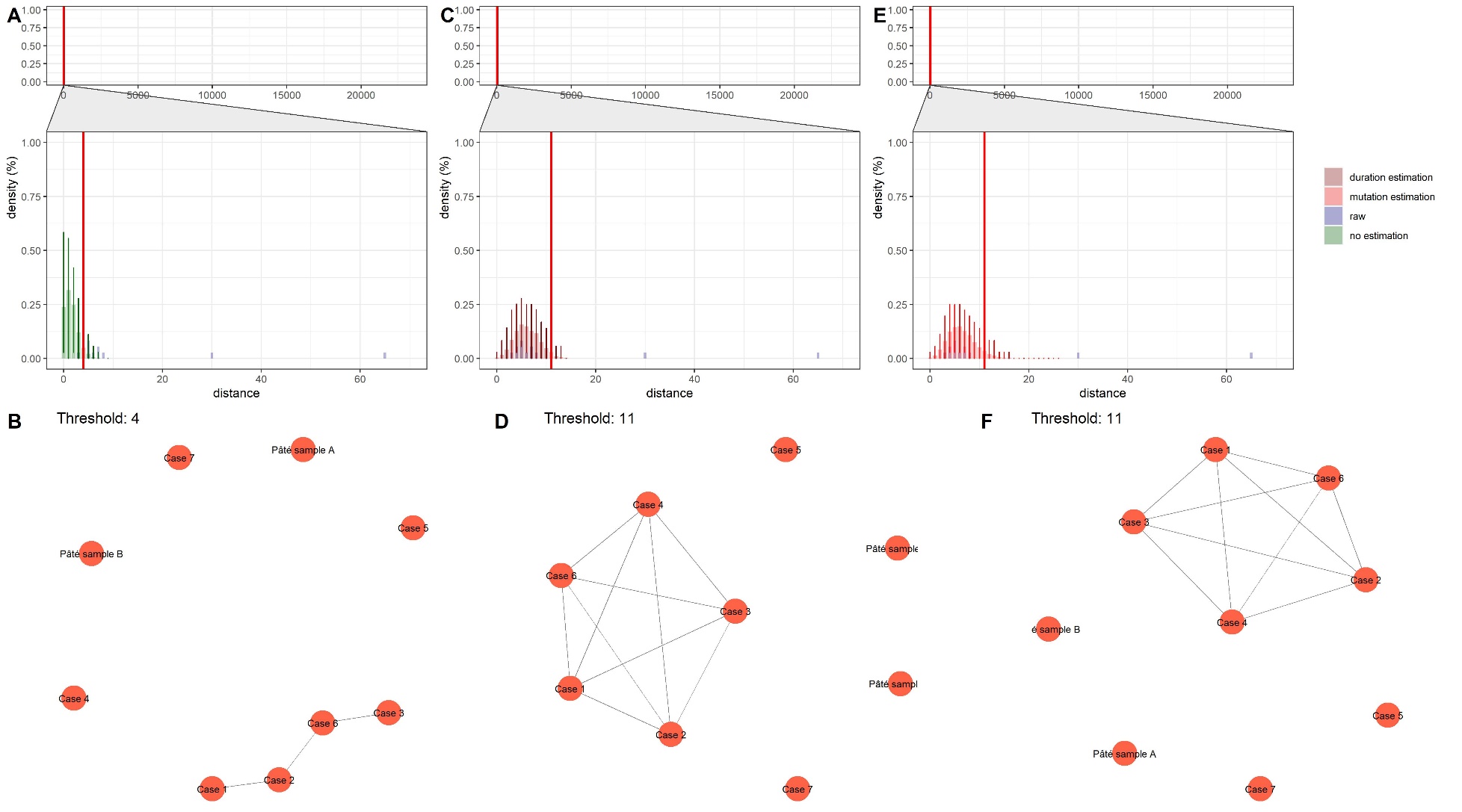

**Figure S4. Analysis of outbreak 4: Comparison between distance thresholds derived from the original publication on the outbreak, and from the current modelling framework.**

In A, C and E are presented the SNP distance distributions: observed (raw) distribution (blue), simulated distribution without estimation (green), simulated with the estimated duration of outbreak (dark red) and simulated with the estimated evolutionary rate (substitution, red). Error bars represent the interval of prediction at 95% of 100 simulations. Red vertical lines correspond to the derived distance threshold. In B, D and F are presented the resulting single-linkage clusters according to the derived distance threshold, defined here as the 99^th^ percentile of the simulated distributions with values corresponding to panels A, C and E, respectively.

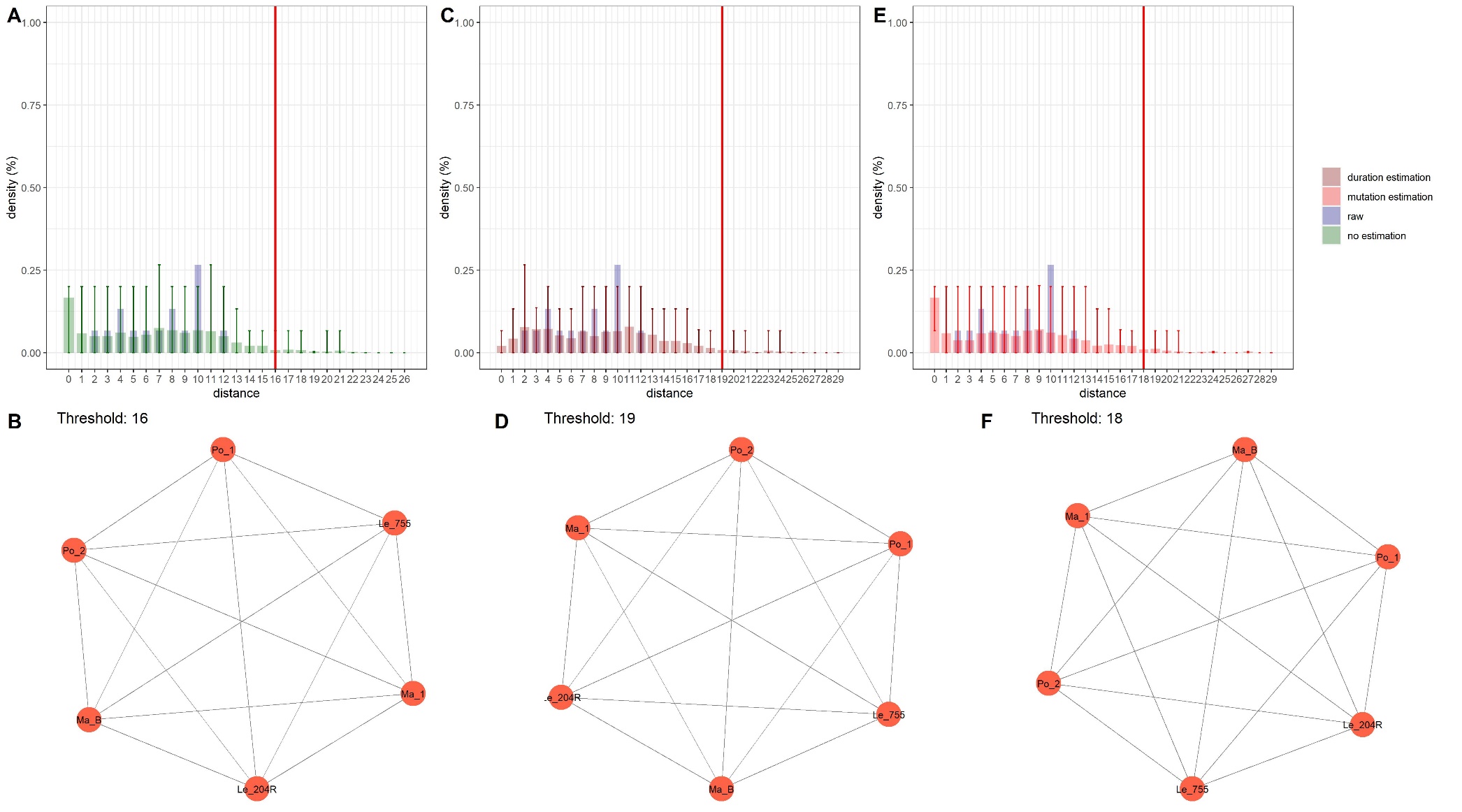

**Figure S5. Analysis of outbreak 5: Comparison between distance thresholds derived from the original publication on the outbreak, and from the current modelling framework.**

In A, C and E are presented the cgMLST distance distributions: observed (raw) distribution (blue), simulated distribution without estimation (green), simulated with the estimated duration of outbreak (dark red) and simulated with the estimated evolutionary rate (substitution, red). Error bars represent the interval of prediction at 95% of 100 simulations. Red vertical lines correspond to the derived distance threshold. In B, D and F are presented the resulting single-linkage clusters according to the derived distance threshold, defined here as the 99^th^ percentile of the simulated distributions with values corresponding to panels A, C and E, respectively.

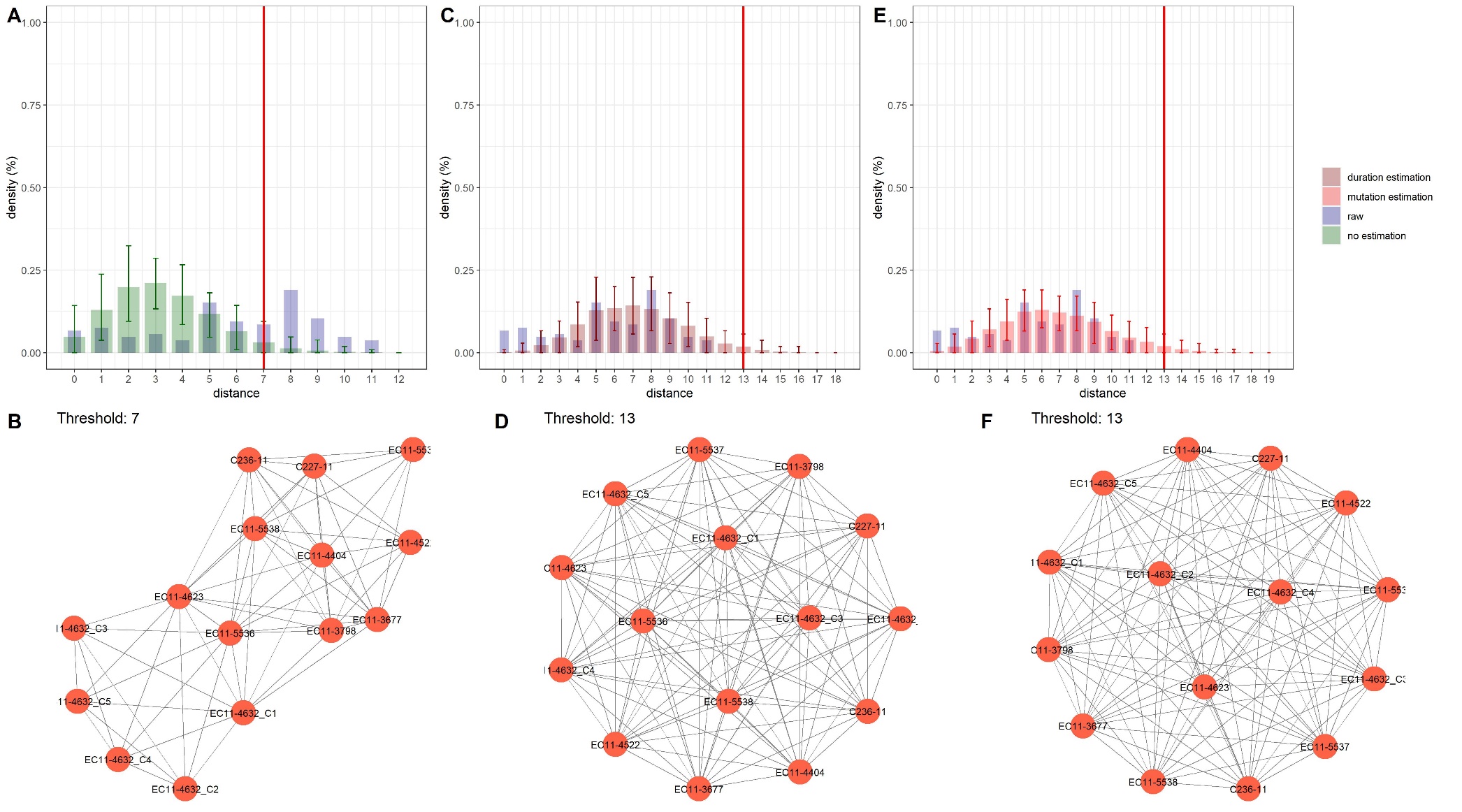

**Figure S6. Analysis of outbreak 6: Comparison between distance thresholds derived from the original publication on the outbreak, and from the current modelling framework.**

In A, C and E are presented the SNP distance distributions: observed (raw) distribution (blue), simulated distribution without estimation (green), simulated with the estimated duration of outbreak (dark red) and simulated with the estimated evolutionary rate (substitution, red). Error bars represent the interval of prediction at 95% of 100 simulations. Red vertical lines correspond to the derived distance threshold. In B, D and F are presented the resulting single-linkage clusters according to the derived distance threshold, defined here as the 99^th^ percentile of the simulated distributions with values corresponding to panels A, C and E, respectively

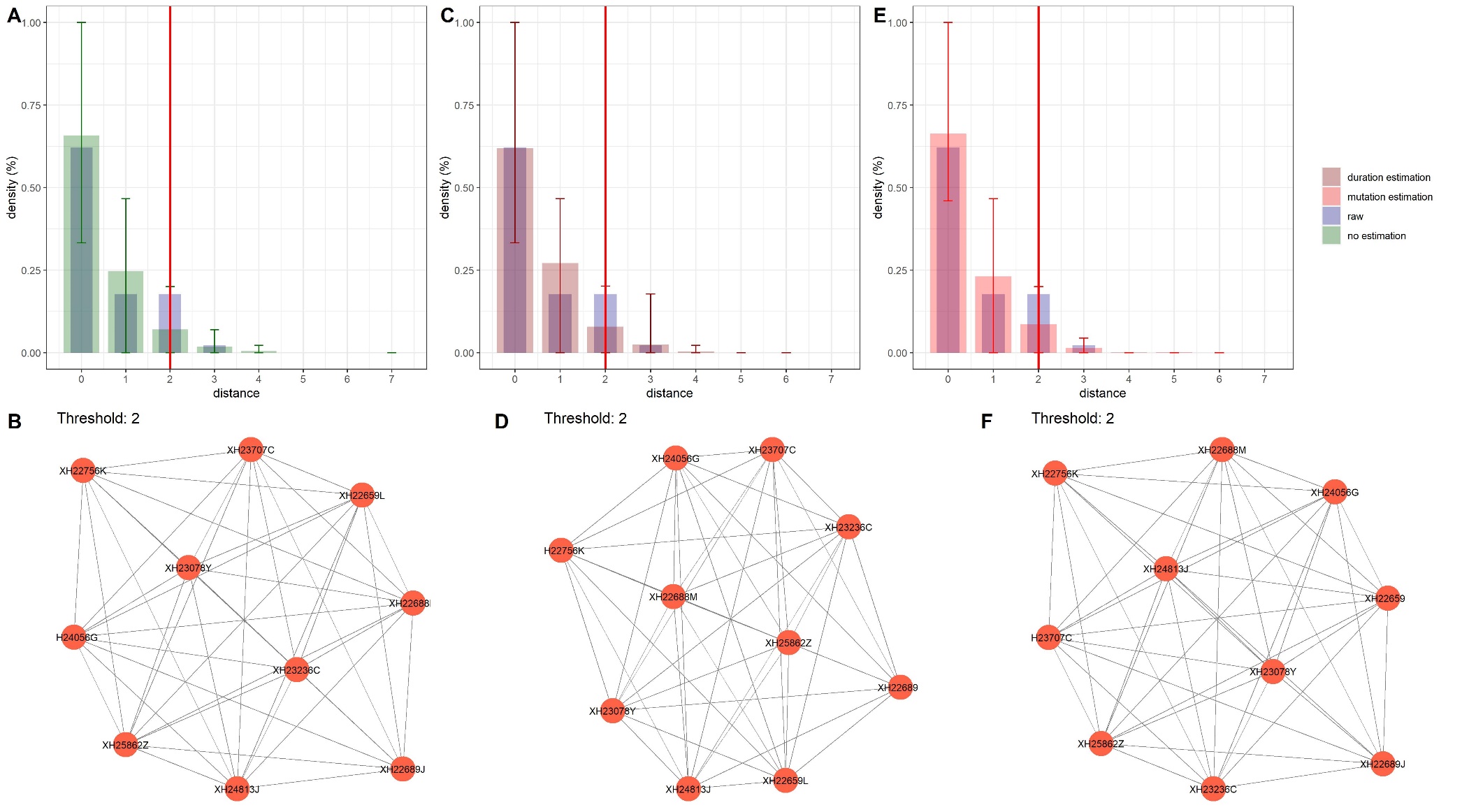

**Figure S7. Analysis of outbreak 7: Comparison between distance thresholds derived from the original publication on the outbreak, and from the current modelling framework.**

In A, C and E are presented the SNP distance distributions: observed (raw) distribution (blue), simulated distribution without estimation (green), simulated with the estimated duration of outbreak (dark red) and simulated with the estimated evolutionary rate (substitution, red). Error bars represent the interval of prediction at 95% of 100 simulations. Red vertical lines correspond to the derived distance threshold. In B, D and F are presented the resulting single-linkage clusters according to the derived distance threshold, defined here as the 99^th^ percentile of the simulated distributions with values corresponding to panels A, C and E, respectively.

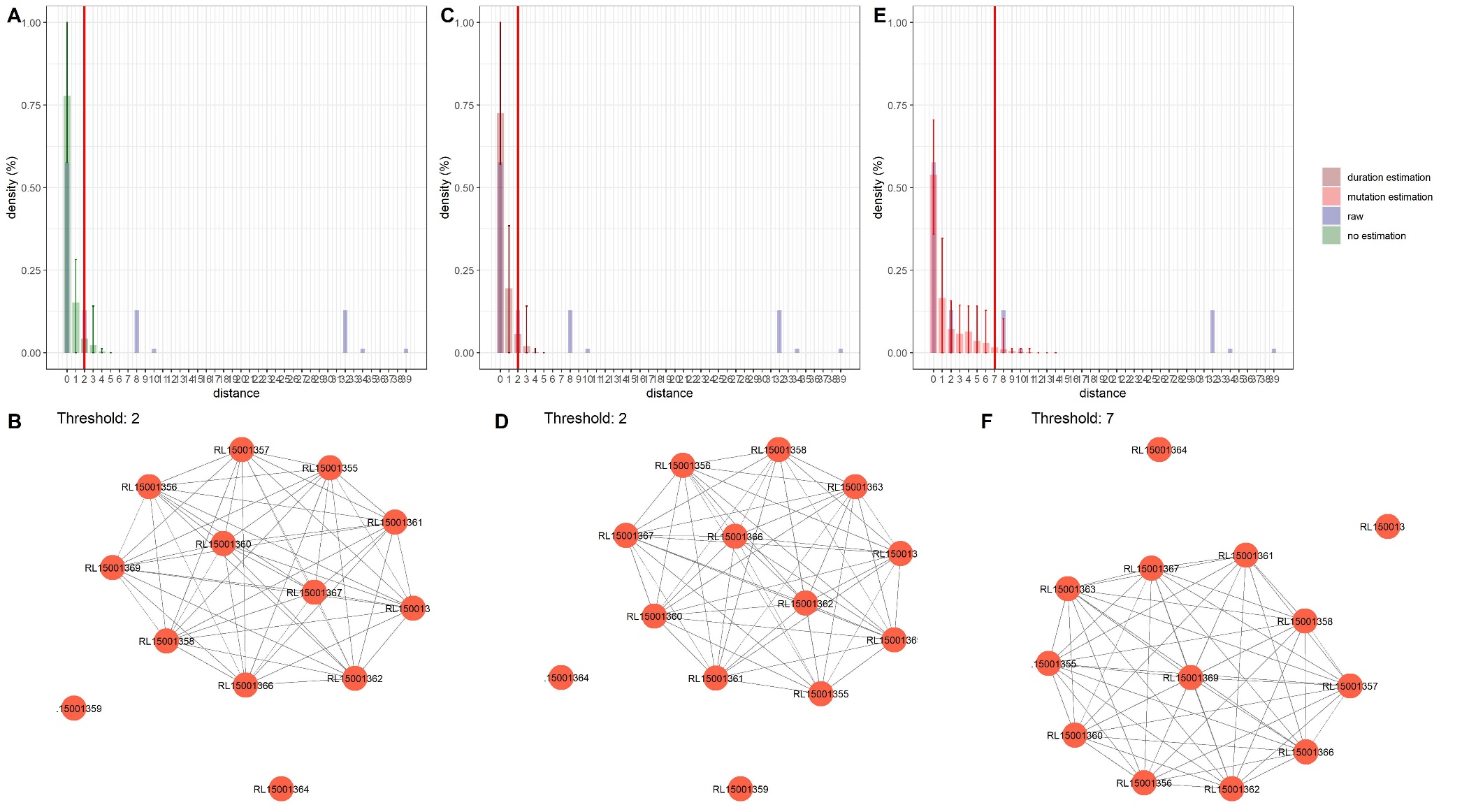

**Figure S8. Analysis of outbreak 8: Comparison between distance thresholds derived from the original publication on the outbreak, and from the current modelling framework.**

In A, C and E are presented the cgMLST distance distributions: observed (raw) distribution (blue), simulated distribution without estimation (green), simulated with the estimated duration of outbreak (dark red) and simulated with the estimated evolutionary rate (substitution, red). Error bars represent the interval of prediction at 95% of 100 simulations. Red vertical lines correspond to the derived distance threshold. In B, D and F are presented the resulting single-linkage clusters according to the derived distance threshold, defined here as the 99^th^ percentile of the simulated distributions with values corresponding to panels A, C and E, respectively.

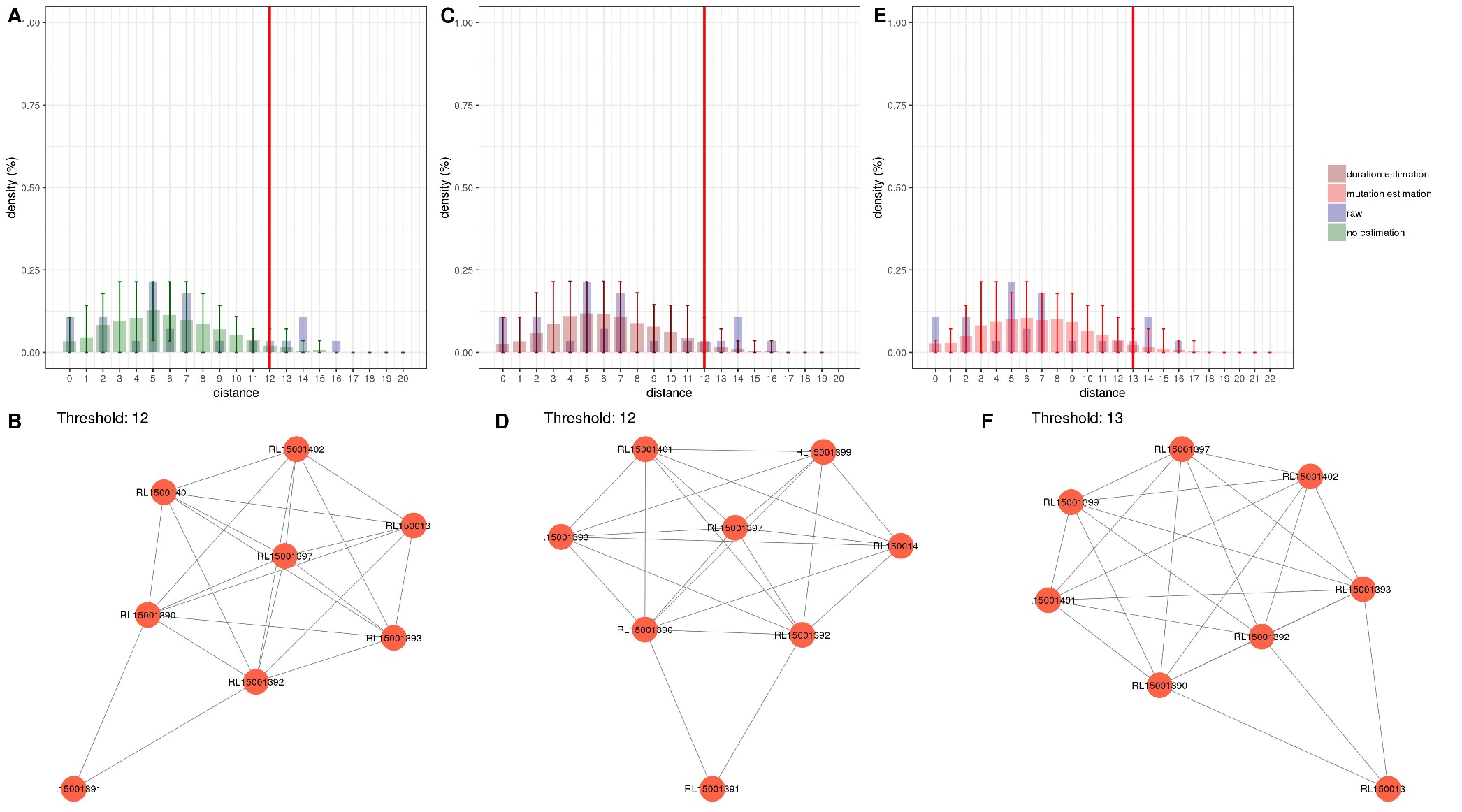

**Figure S9. Analysis of outbreak 9: Comparison between distance thresholds derived from the original publication on the outbreak, and from the current modelling framework.**

In A, C and E are presented the cgMLST distance distributions: observed (raw) distribution (blue), simulated distribution without estimation (green), simulated with the estimated duration of outbreak (dark red) and simulated with the estimated evolutionary rate (substitution, red). Error bars represent the interval of prediction at 95% of 100 simulations. Red vertical lines correspond to the derived distance threshold. In B, D and F are presented the resulting single-linkage clusters according to the derived distance threshold, defined here as the 99^th^ percentile of the simulated distributions with values corresponding to panels A, C and E, respectively.

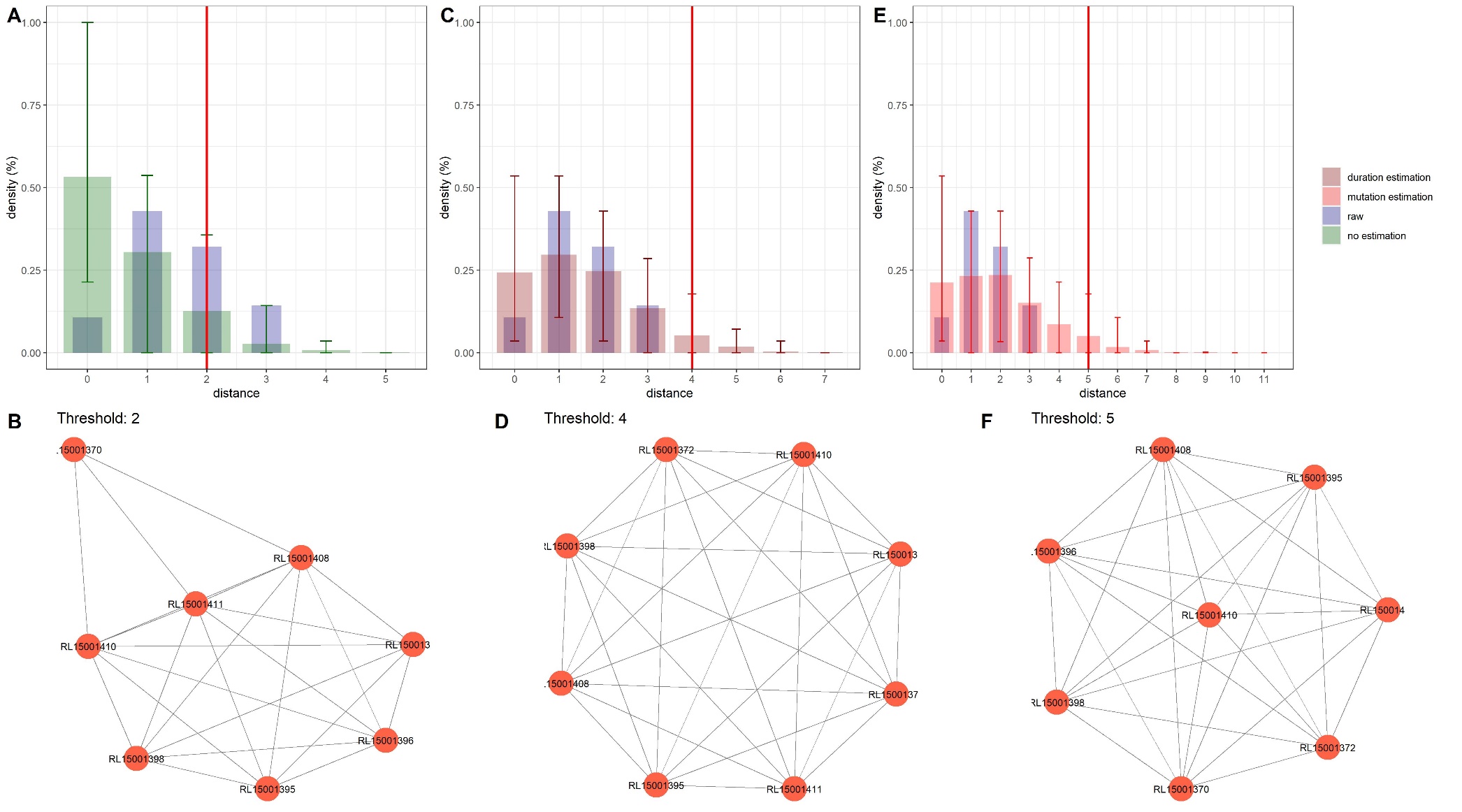

**Figure S10. Analysis of outbreak 10: Comparison between distance thresholds derived from the original publication on the outbreak, and from the current modelling framework.**

In A, C and E are presented the cgMLST distance distributions: observed (raw) distribution (blue), simulated distribution without estimation (green), simulated with the estimated duration of outbreak (dark red) and simulated with the estimated evolutionary rate (substitution, red). Error bars represent the interval of prediction at 95% of 100 simulations. Red vertical lines correspond to the derived distance threshold. In B, D and F are presented the resulting single-linkage clusters according to the derived distance threshold, defined here as the 99^th^ percentile of the simulated distributions with values corresponding to panels A, C and E, respectively.

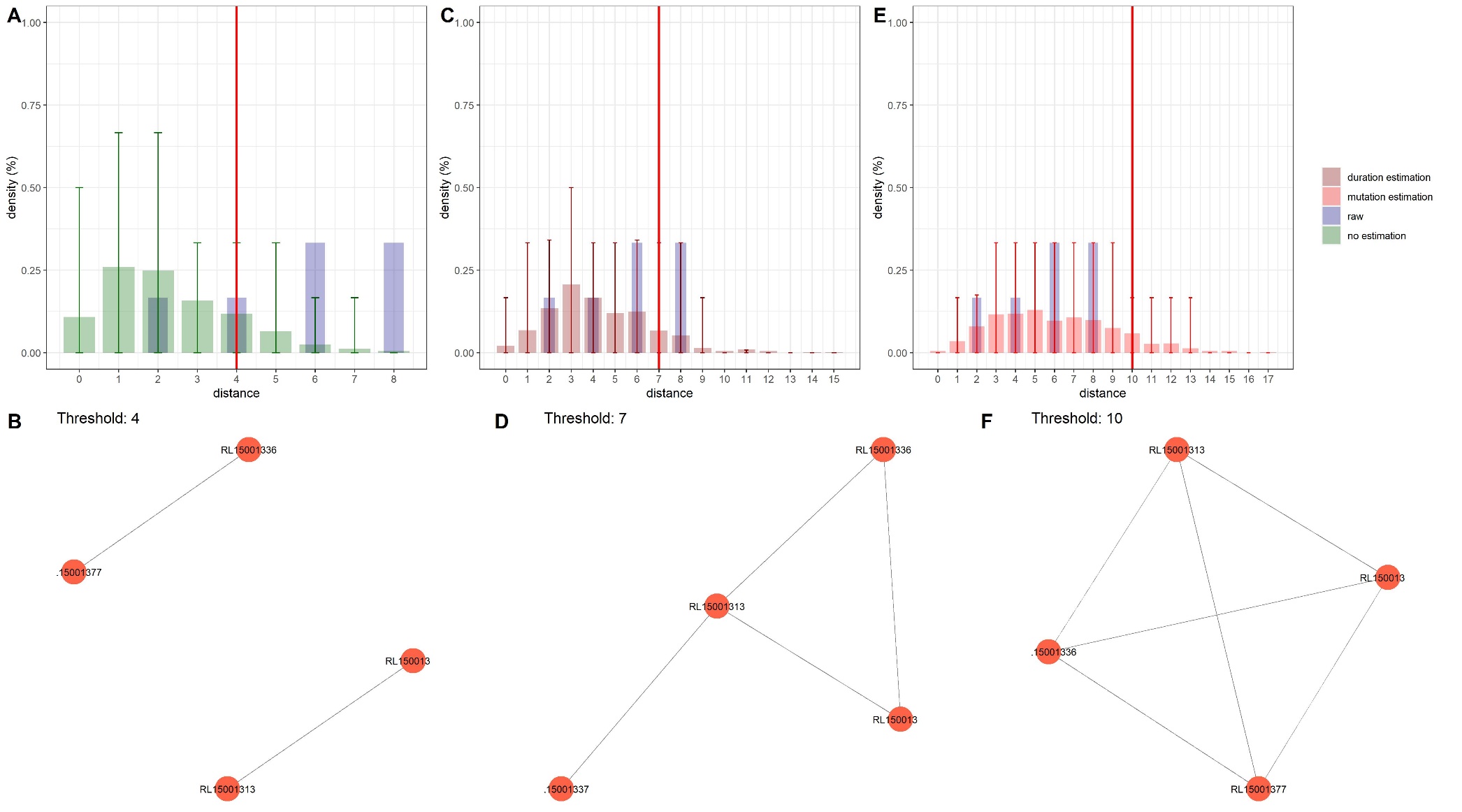

**Figure S11. Analysis of outbreak 11: Comparison between distance thresholds derived from the original publication on the outbreak, and from the current modelling framework.**

In A, C and E are presented the cgMLST distance distributions: observed (raw) distribution (blue), simulated distribution without estimation (green), simulated with the estimated duration of outbreak (dark red) and simulated with the estimated evolutionary rate (substitution, red). Error bars represent the interval of prediction at 95% of 100 simulations. Red vertical lines correspond to the derived distance threshold. In B, D and F are presented the resulting single-linkage clusters according to the derived distance threshold, defined here as the 99^th^ percentile of the simulated distributions with values corresponding to panels A, C and E, respectively.

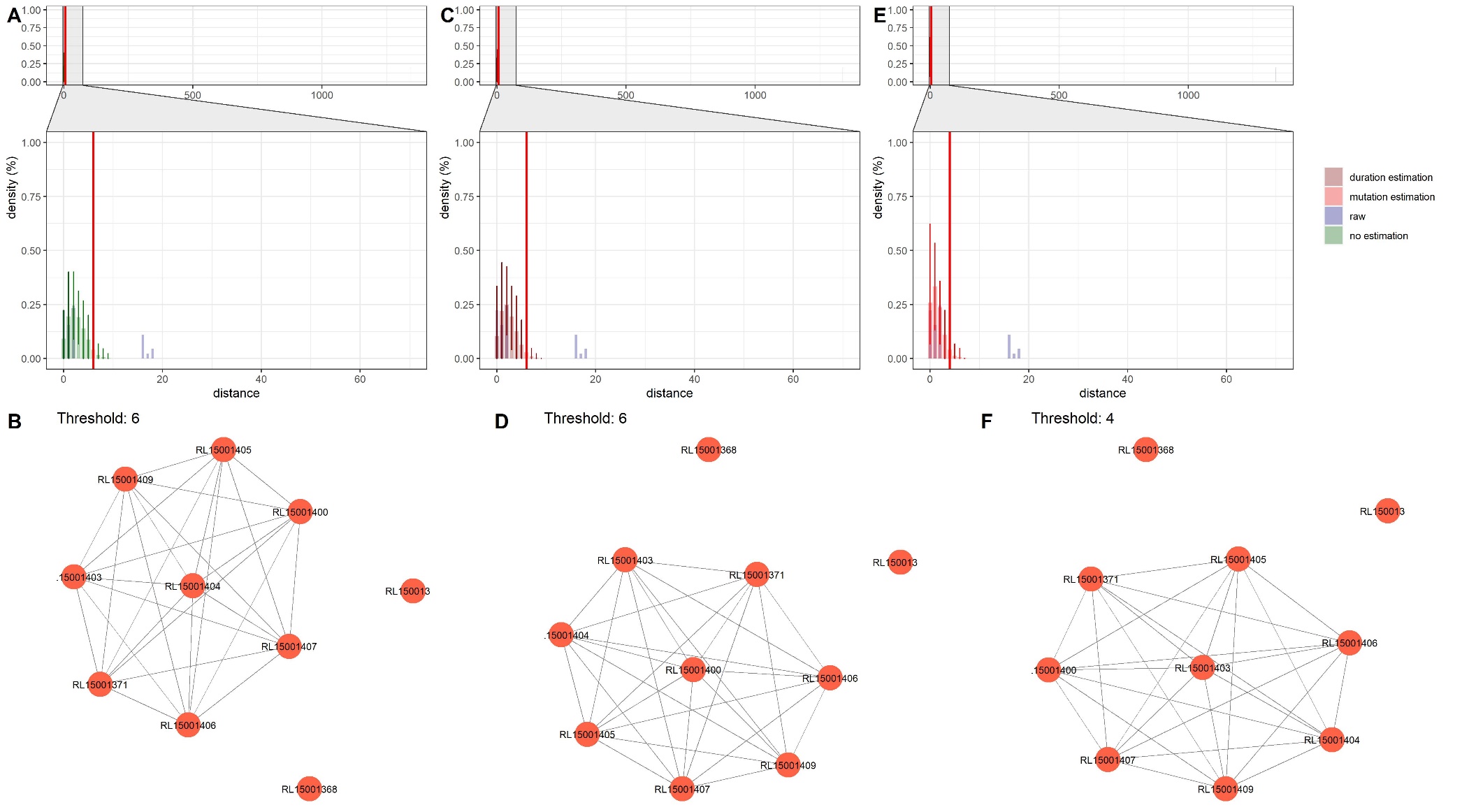

**Figure S12. Analysis of outbreak 12: Comparison between distance thresholds derived from the original publication on the outbreak, and from the current modelling framework.**

In A, C and E are presented the cgMLST distance distributions: observed (raw) distribution (blue), simulated distribution without estimation (green), simulated with the estimated duration of outbreak (dark red) and simulated with the estimated evolutionary rate (substitution, red). Error bars represent the interval of prediction at 95% of 100 simulations. Red vertical lines correspond to the derived distance threshold. In B, D and F are presented the resulting single-linkage clusters according to the derived distance threshold, defined here as the 99^th^ percentile of the simulated distributions with values corresponding to panels A, C and E, respectively.

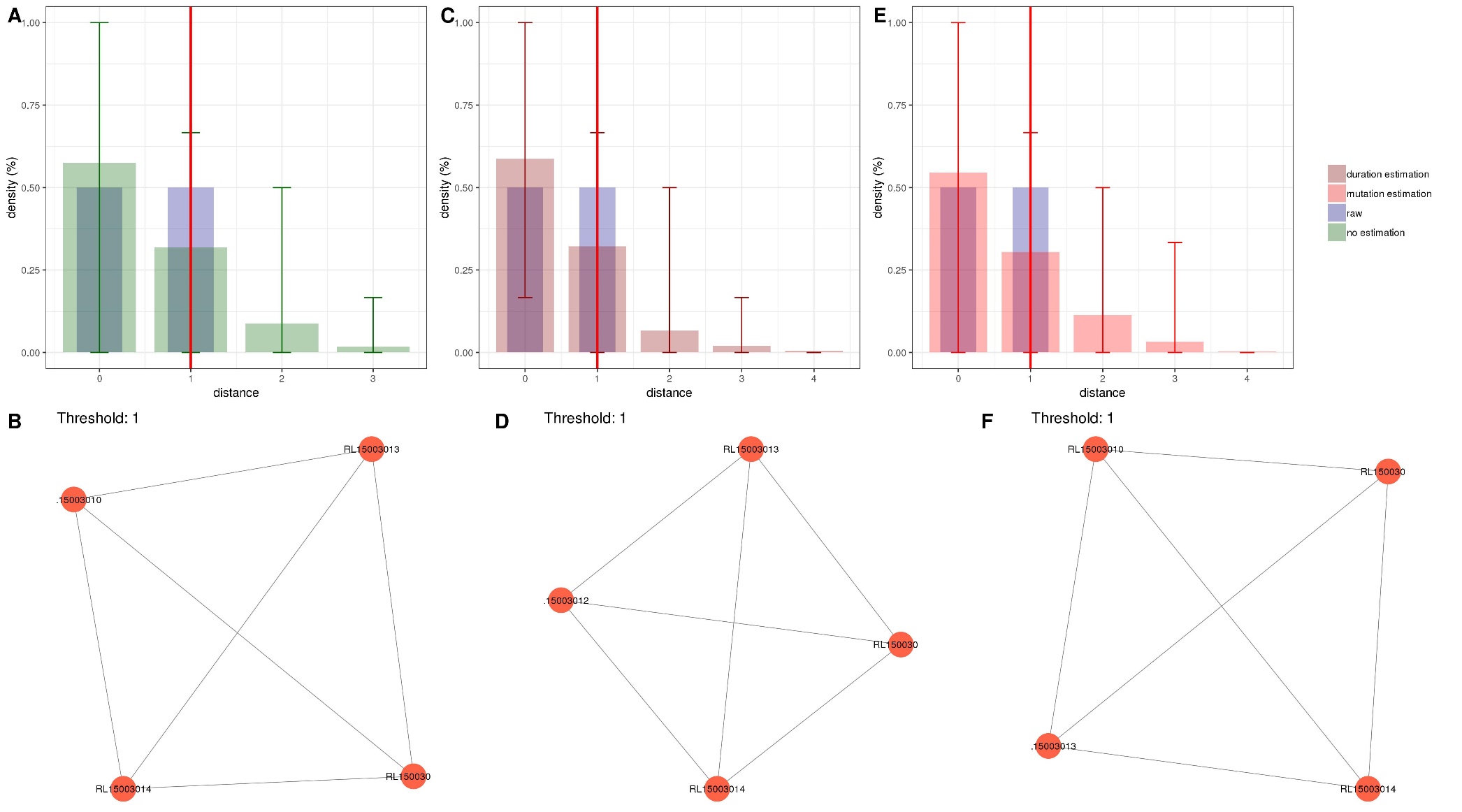

**Figure S13. Analysis of outbreak 13: Comparison between distance thresholds derived from the original publication on the outbreak, and from the current modelling framework.**

In A, C and E are presented the cgMLST distance distributions: observed (raw) distribution (blue), simulated distribution without estimation (green), simulated with the estimated duration of outbreak (dark red) and simulated with the estimated evolutionary rate (substitution, red). Error bars represent the interval of prediction at 95% of 100 simulations. Red vertical lines correspond to the derived distance threshold. In B, D and F are presented the resulting single-linkage clusters according to the derived distance threshold, defined here as the 99^th^ percentile of the simulated distributions with values corresponding to panels A, C and E, respectively.

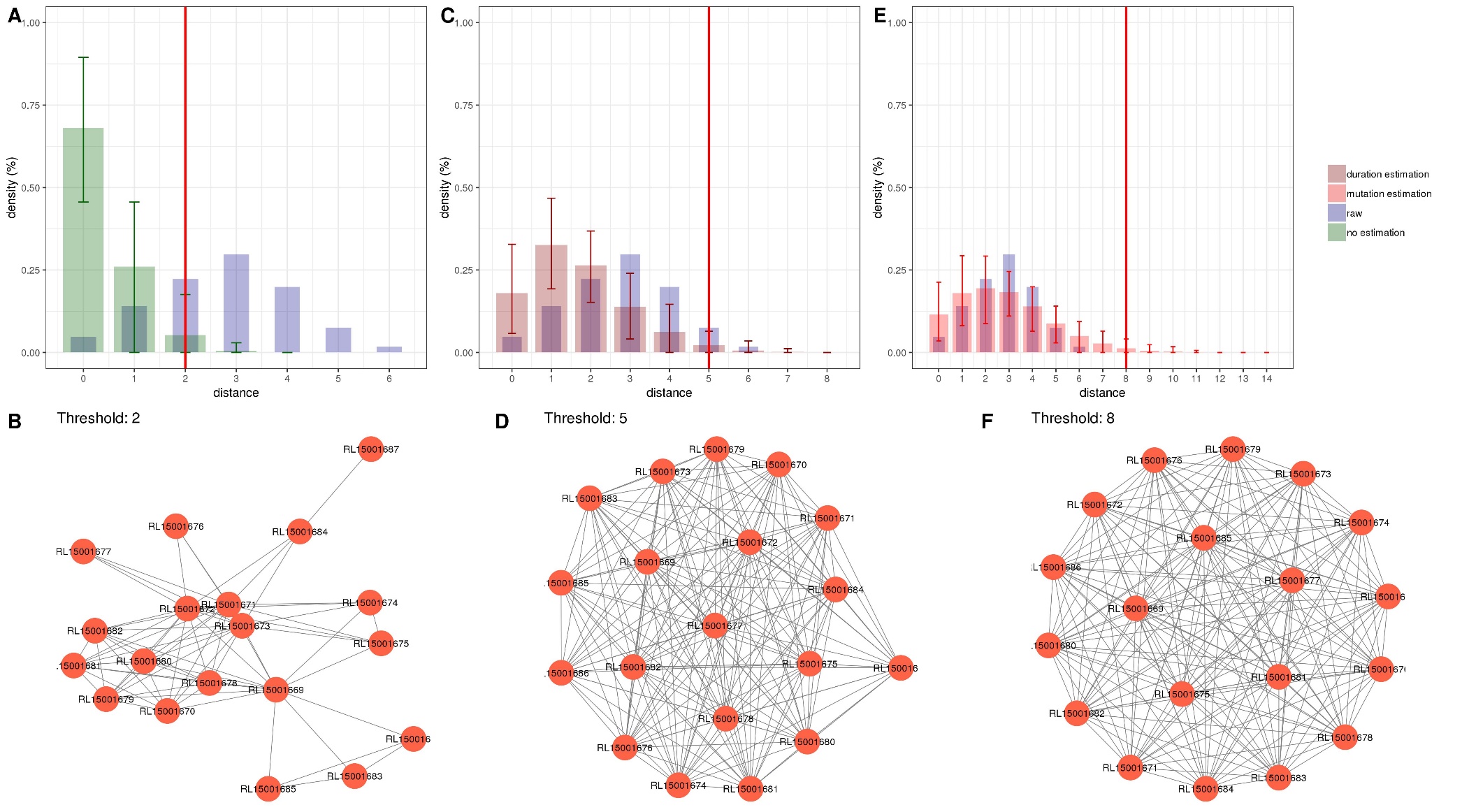

**Figure S14. Analysis of outbreak 14: Comparison between distance thresholds derived from the original publication on the outbreak, and from the current modelling framework.**

In A, C and E are presented the cgMLST distance distributions: observed (raw) distribution (blue), simulated distribution without estimation (green), simulated with the estimated duration of outbreak (dark red) and simulated with the estimated evolutionary rate (substitution, red). Error bars represent the interval of prediction at 95% of 100 simulations. Red vertical lines correspond to the derived distance threshold. In B, D and F are presented the resulting single-linkage clusters according to the derived distance threshold, defined here as the 99^th^ percentile of the simulated distributions with values corresponding to panels A, C and E, respectively.

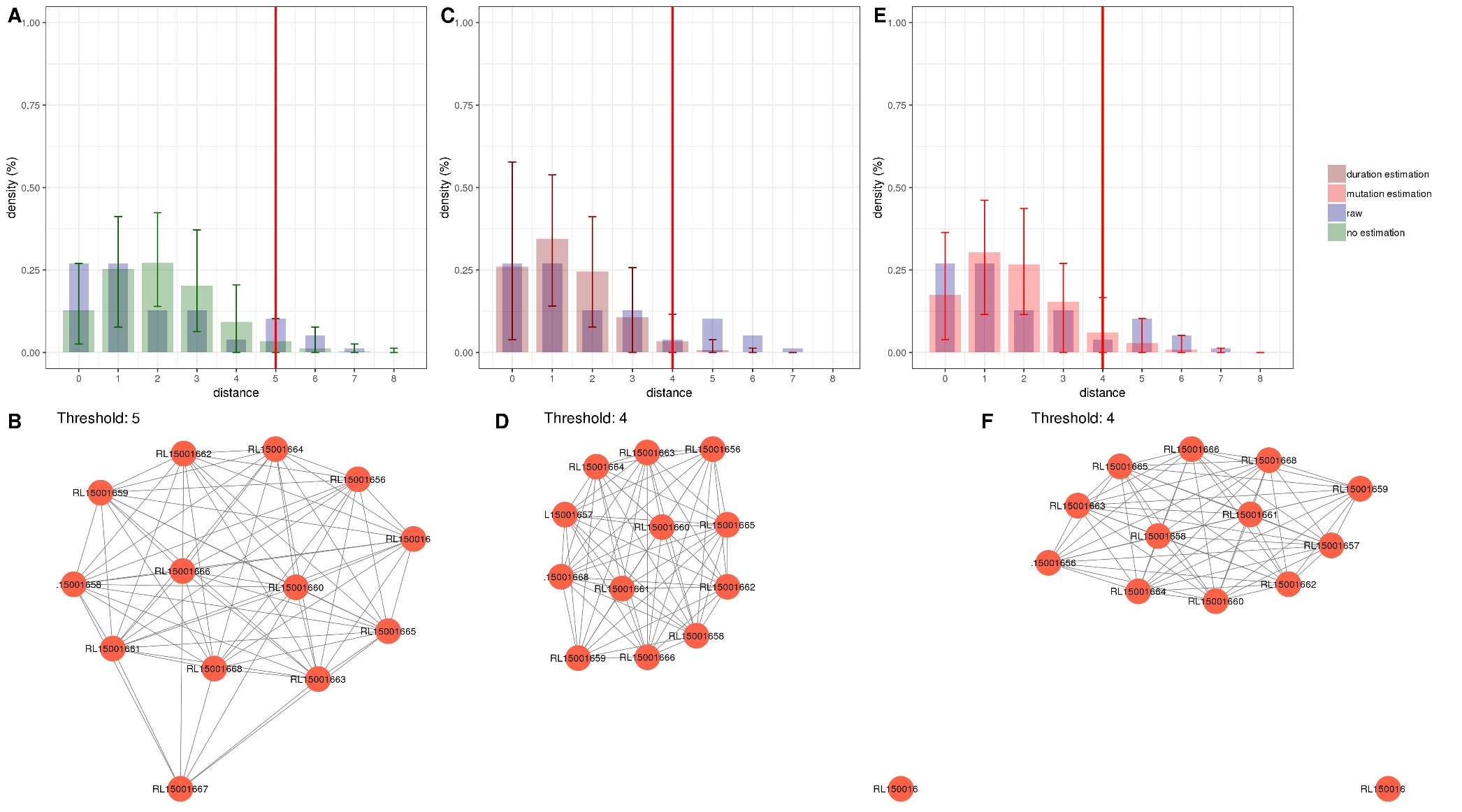

**Figure S15. Analysis of outbreak 15: Comparison between distance thresholds derived from the original publication on the outbreak, and from the current modelling framework.**

In A, C and E are presented the cgMLST distance distributions: observed (raw) distribution (blue), simulated distribution without estimation (green), simulated with the estimated duration of outbreak (dark red) and simulated with the estimated evolutionary rate (substitution, red). Error bars represent the interval of prediction at 95% of 100 simulations. Red vertical lines correspond to the derived distance threshold. In B, D and F are presented the resulting single-linkage clusters according to the derived distance threshold, defined here as the 99^th^ percentile of the simulated distributions with values corresponding to panels A, C and E, respectively.

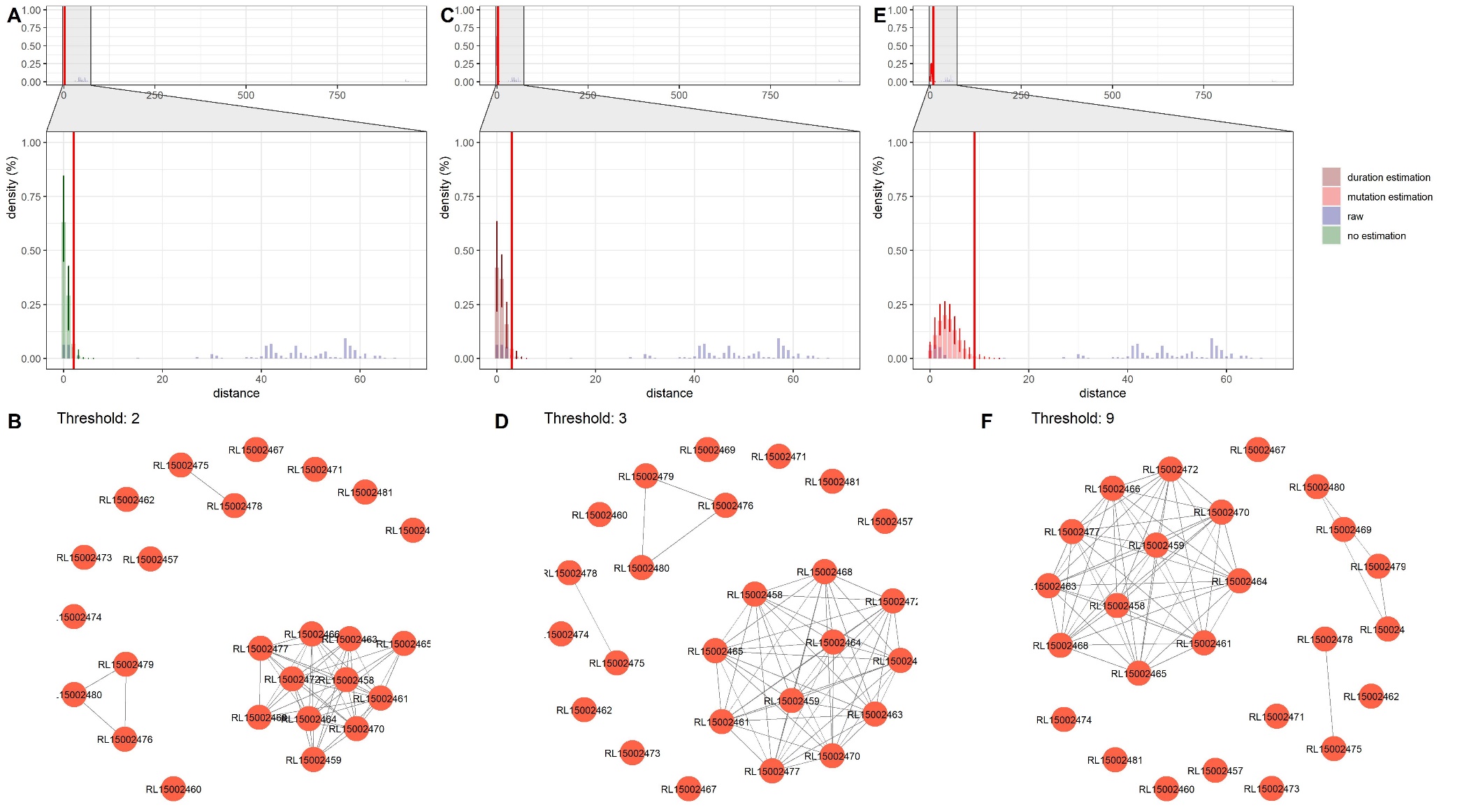

**Figure S16. Analysis of outbreak 16: Comparison between distance thresholds derived from the original publication on the outbreak, and from the current modelling framework.**

In A, C and E are presented the cgMLST distance distributions: observed (raw) distribution (blue), simulated distribution without estimation (green), simulated with the estimated duration of outbreak (dark red) and simulated with the estimated evolutionary rate (substitution, red). Error bars represent the interval of prediction at 95% of 100 simulations. Red vertical lines correspond to the derived distance threshold. In B, D and F are presented the resulting single-linkage clusters according to the derived distance threshold, defined here as the 99^th^ percentile of the simulated distributions with values corresponding to panels A, C and E, respectively.

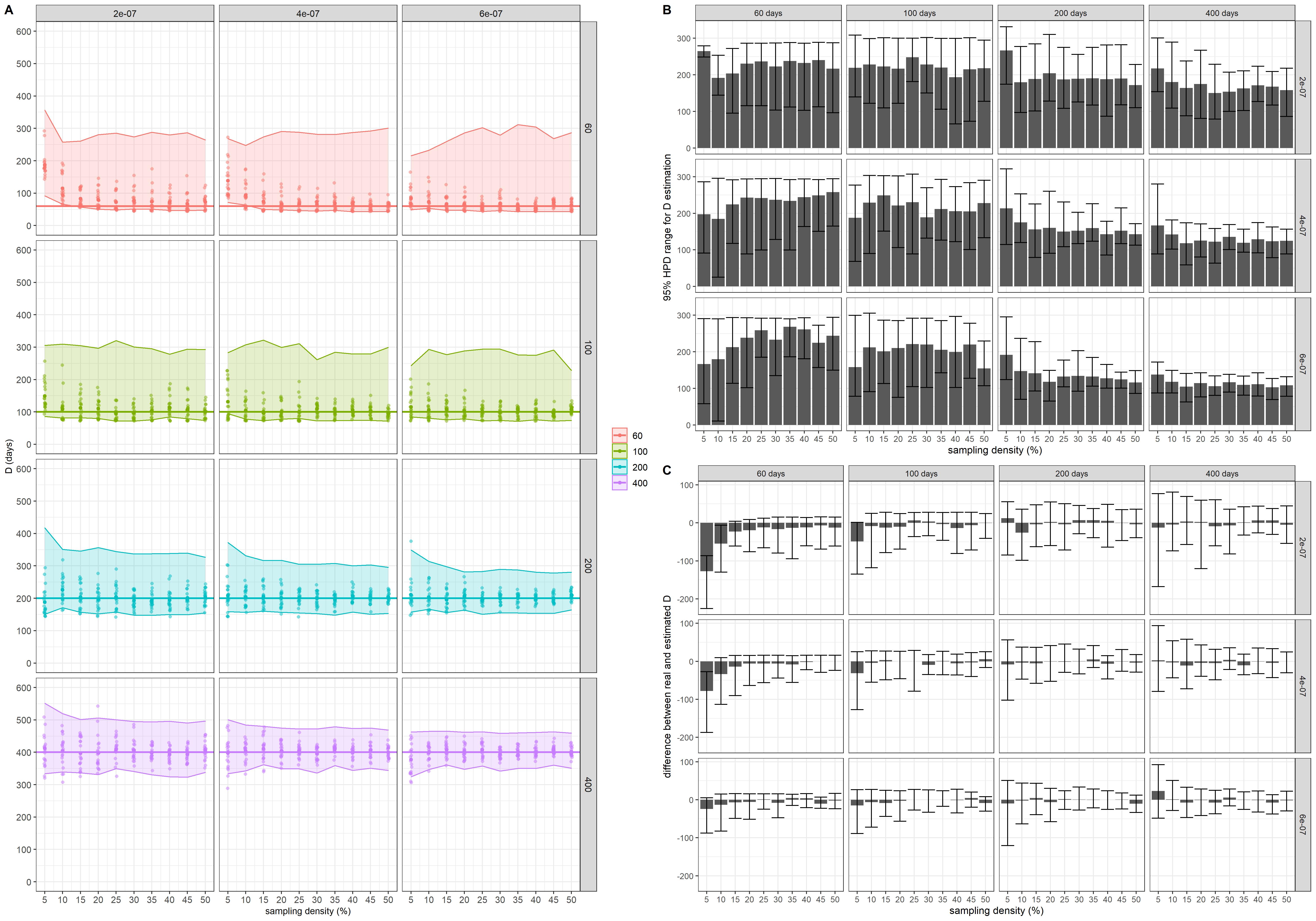

**Figure S17. D estimation analysis.** In A, average best estimation of *D* of the three MCMC chains for each of the 20 outbreaks. In B, average HPD range of the 20 outbreaks for 10 sampling densities (from 5% to 50%). In C, average distance between real and best estimated *D*. In B and C, error bars correspond to the 95% prediction interval.

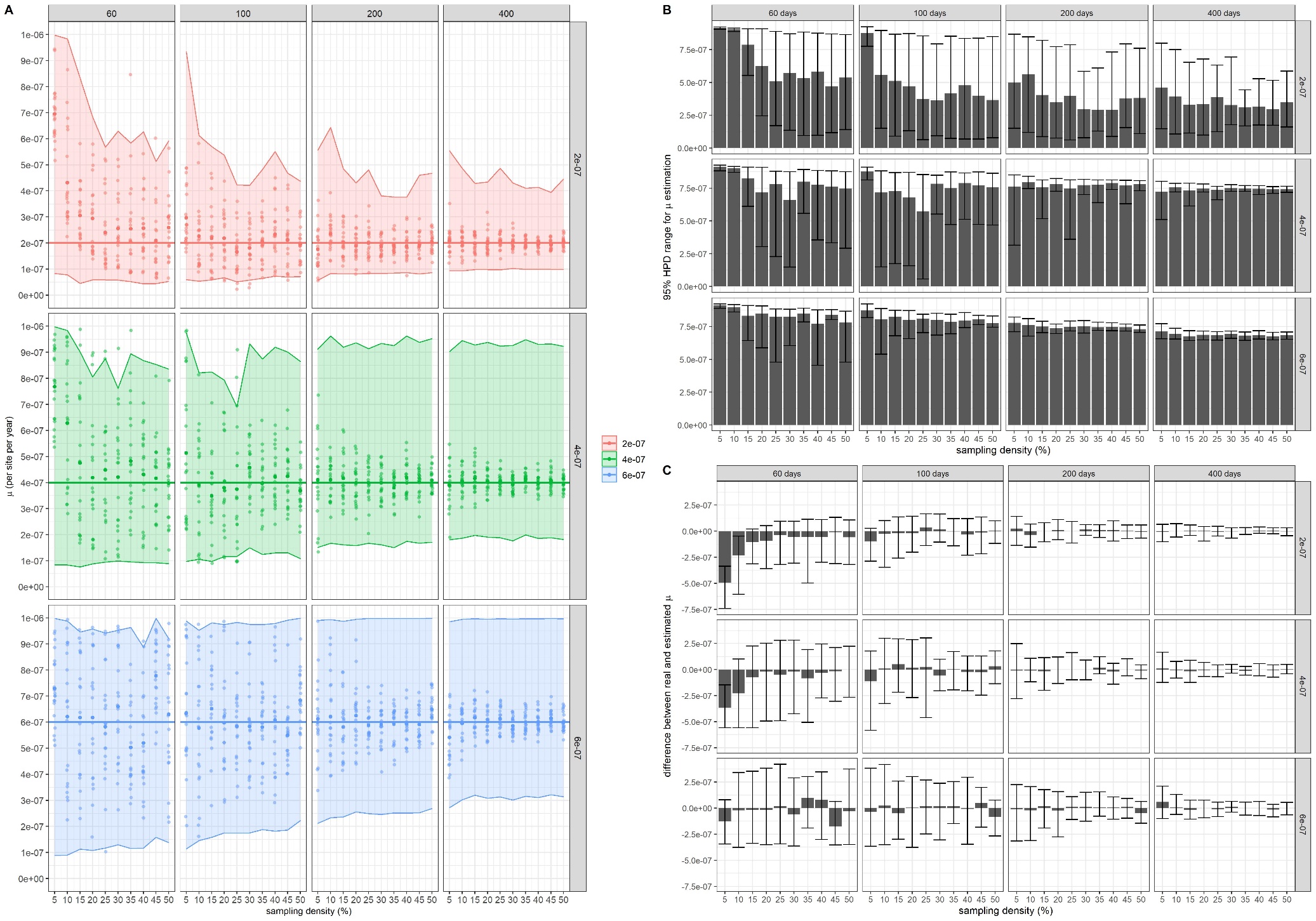

**Figure S18. *μ* estimation analysis.** In A, average best estimation of *μ* of the three MCMC chains for each of the 20 outbreaks. In B, average HPD range of the 20 outbreaks for 10 sampling densities (5 to 50%). In C, average distance between real and best estimated *μ*. In B and C, error bars correspond to the 95% prediction interval.
